# Transcranial magnetic stimulation to the dorsolateral prefrontal cortex modulates single-neuron activity in humans

**DOI:** 10.64898/2026.03.15.711839

**Authors:** Charles W. Dickey, Umair Hassan, Hiroto Kawasaki, Ariane E. Rhone, Christopher C. Cline, Matthew A. Howard, Nicholas T. Trapp, Aaron D. Boes, Joel I. Berger, Corey J. Keller

**Author notes:** Co-senior authors (Aaron D. Boes, Joel I. Berger, Corey J. Keller). **Author Contributions:** Charles W. Dickey: Conceptualization, Methodology, Formal analysis, Visualization, Writing - original draft, Writing - review & editing. Umair Hassan: Data curation, Visualization, Writing - review & editing. Hiroto Kawasaki: Investigation. Ariane E. Rhone: Investigation, Writing - review & editing. Christopher C. Cline: Investigation, Writing - review & editing. Matthew A. Howard: Resources, Funding acquisition. Nicholas T. Trapp: Investigation, Writing - review & editing. Aaron D. Boes: Conceptualization, Investigation, Supervision, Writing - review & editing, Funding acquisition. Joel I. Berger: Conceptualization, Investigation, Formal analysis, Supervision, Writing - review & editing, Funding acquisition. Corey J. Keller: Conceptualization, Supervision, Writing - review & editing, Funding acquisition.

## Abstract

Transcranial magnetic stimulation (TMS) to the dorsolateral prefrontal cortex (dlPFC) is an FDA-cleared treatment for depression, yet how cortical stimulation influences single neurons in deep brain circuits remains unknown. Using intracranial microelectrode recordings in four neurosurgical patients, we resolved single-neuron spikes as early as 8 ms from 185 single neurons after single-pulse left dlPFC TMS. TMS elicited time-locked firing responses in 46% of neurons across deep cortical and subcortical structures bilaterally. TMS facilitated putative interneuron spiking in striato-thalamic regions from ∼8 ms, peaking at ∼80-100 ms, and lasting to ∼1000 ms, while suppressing putative pyramidal cell spiking with a delayed and slower time course. Trial-by-trial single-neuron modulations were positively correlated with cortico-striato-thalamic network activity and anti-correlated with limbic network activity. These findings reveal that dlPFC TMS facilitates inhibitory firing in executive control networks while suppressing limbic excitatory drive, providing a cellular mechanism for how cortical stimulation modulates distributed brain networks.

## Introduction

Transcranial magnetic stimulation (TMS) to the dorsolateral prefrontal cortex (dlPFC) is an FDA-cleared treatment for major depressive disorder (O’Reardon et al., 2007), yet the precise neural mechanisms underlying its therapeutic effects remain debated. The prevailing hypothesis posits that TMS works through engagement of cortico-limbic pathways, particularly by modulating the network connecting the dlPFC to deeper limbic structures including the subgenual anterior cingulate cortex and amygdala (Eshel et al., 2020; Fox et al., 2012; Oathes et al., 2021). According to this model, repeated activation of this pathway induces synaptic plasticity and circuit reorganization that normalizes hyperactive limbic regions, leading to sustained symptom relief.

However, emerging network-level models suggest an alternative mechanism: rather than directly dampening limbic hyperactivity, TMS may restore therapeutic balance by engaging associative cortico-striato-thalamic loops that provide top-down executive control over affective systems (Drysdale et al., 2017; Sheline et al., 2010). These loops - which involve prefrontal cortex, striatum (basal ganglia), and thalamus - regulate cognitive flexibility, action selection, and inhibitory control over limbic reactivity. fMRI and intracranial EEG studies support this hypothesis by demonstrating that dlPFC stimulation modulates activity in these deep striatal and thalamic structures (Dowdle et al., 2018; Solomon et al., 2024; Wang et al., 2024). Importantly, these two mechanisms imply fundamentally different therapeutic actions: one suggesting direct normalization of limbic circuits, the other suggesting rebalancing of large-scale control networks through striato-thalamic gating. While recent intracranial evidence in two participants shows that single-pulse TMS to the dlPFC suppresses high-frequency activity in the subgenual anterior cingulate (Solomon et al., 2026), a major node in the limbic network, it remains unclear whether this represents direct cortico-limbic inhibition or indirect modulation through intervening network nodes. Distinguishing between these possibilities requires direct single-neuron recordings across distributed brain regions, which to date have not been possible in humans (Tsang et al., 2025).

Animal studies have provided initial evidence that TMS can modulate cortical neurons, typically inducing early facilitation followed by suppression, though these recordings have been largely restricted to the local site of stimulation (Allen et al., 2007; Moliadze et al., 2003; Mueller et al., 2014; Romero et al., 2019; Vlachos et al., 2012). A knowledge gap exists regarding how stimulation affects single neurons in deeper, distributed brain regions thought to mediate therapeutic effects in humans. To our knowledge, only one study has measured single-neuron responses to TMS in humans, where in patients with Parkinson’s disease, motor cortex stimulation modulated activity in the ipsilateral subthalamic nucleus (Strafella et al., 2004). No study has investigated the neuronal effects of TMS to the dlPFC, which is the primary target for depression treatment. We hypothesized that single-pulse dlPFC TMS directly recruits inhibitory striato-thalamic (executive control) neurons and indirectly suppresses excitatory limbic neurons.

In the first investigation of human single-neuron responses to dlPFC TMS, we found that stimulation facilitated spiking within striato-thalamic regions while suppressing spiking in limbic regions. Across 185 single units recorded from four participants, 27% showed increased firing, concentrated from ∼8-300 ms, whereas 21% exhibited suppression. These effects were observed bilaterally and facilitation largely localized to striato-thalamic neurons whereas suppression preferentially localized to limbic neurons. Cellular classification revealed that putative interneurons in striato-thalamic areas were facilitated whereas putative pyramidal cells were suppressed in widespread areas. Concurrent macroelectrode recordings revealed that trial-by-trial spiking modulations were positively correlated with cortico-striato-thalamic network activity and anti-correlated with limbic network activity. Together, these findings show that left dlPFC TMS selectively recruits inhibitory neurons in executive and thalamic relay circuits while dampening excitatory limbic drive, revealing how prefrontal stimulation propagates through cortico-subcortical circuits.

## Results

### Single-unit spikes resolved ∼8 ms after dlPFC TMS in humans

We delivered single-pulse TMS to the left dlPFC in four participants (3 female, ages 35-56; Supplementary Table 1) with medication-resistant epilepsy who were implanted with intracranial depth electrodes for seizure focus localization (see Methods for details). TMS was targeted to dlPFC using the modified Beam F3 approach (Beam et al., 2009; Trapp et al., 2020) and was delivered solely for the purpose of research. TMS was delivered toward the end of the clinical monitoring period, which was 12-19 days after electrode implantation (Supplementary Table 1). Each participant received blocks of 50 single pulses of active TMS. In addition, sham TMS blocks were delivered with the TMS coil flipped 180° to generate the acoustic stimulus while minimizing brain stimulation. In this configuration, the induced electromagnetic field was directed away from the head and shielded with a nickel-iron alloy (Figure 1A). Across the four participants, 58 depth electrodes (range=14-15) were implanted, of which 37 of these (range=7-11) had bundles of eight platinum-iridium microwires extending 2-4 mm from the electrode tip, referenced to a ninth uninsulated wire within each bundle (Figure 1B). Microwires were distributed across deep cortical and subcortical structures bilaterally, including the putamen, thalamus, hippocampus, amygdala, and multiple divisions of the cingulate cortex (Figure 1C, Supplementary Table 1).

**Figure 1:**
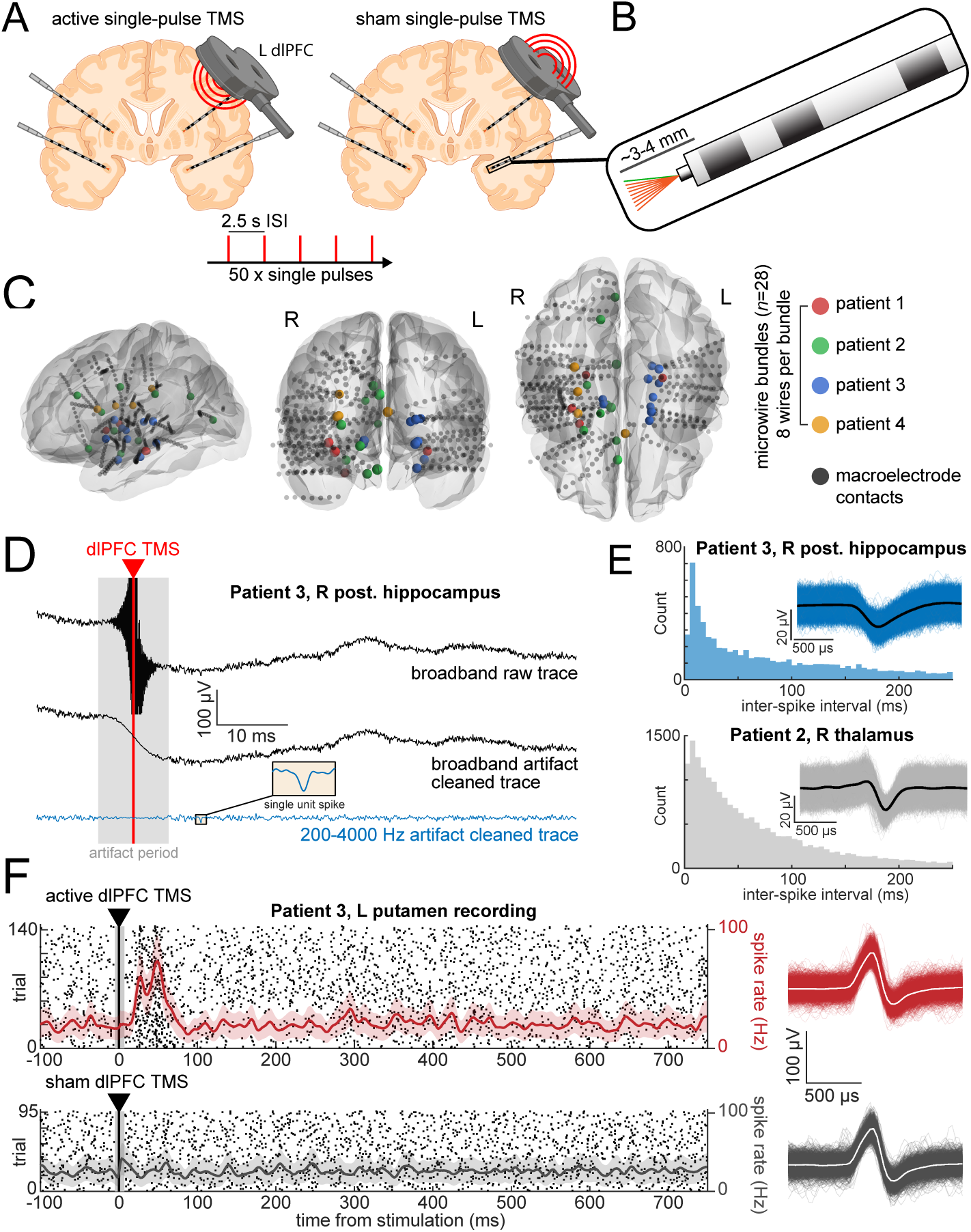
Human single-unit activity recorded in response to single-pulse TMS to the left dlPFC. (A) Active and sham single-pulse TMS was delivered at 100-120% resting motor threshold to the left dlPFC in participants implanted with intracranial EEG Behnke-Fried depth electrodes.Microwires extending from the depth electrode tips enabled single-unit recordings. Multiple blocks of 50 single pulses were delivered with a 2.5-3 s inter-stimulus interval. Sham TMS was delivered by flipping the TMS coil 180° from the active condition. Schematic produced in BioRender. (B) Microwire bundle configuration showing eight insulated recording microwires in orange referenced to a ninth uninsulated reference wire in green extending 3-4 mm from the depth electrode tip. Macroelectrode contacts are shown in dark gray. (C) Anatomical localizations of recording sites across all four participants displayed on fsaverage template brains. Colored spheres indicate microelectrode bundle sites where single units were recorded. Gray spheres show all macroelectrode contact localizations within the brain. (D) Example electrophysiological recordings from a single microwire in the right posterior hippocampus showing signal processing steps. Top shows the broadband raw trace (TMS artifact is truncated in amplitude dimension for visualization). Middle shows the broadband artifact cleaned trace. Bottom trace in blue shows the 200-4000 Hz bandpass filtered artifact cleaned trace used for spike detection, with an isolated single-unit spike waveform shown in the inset. Gray shading indicates the artifact period. See further details on artifact detection in Supplementary Figure 1. (E) Single-unit inter-spike interval histograms and superimposed spike waveforms (individual spikes in color or gray and mean in black). Top panel data correspond to the unit detected with a spike shown in the blue trace in D. Bottom shows data from the right thalamus. Single-unit quality and isolation was confirmed using standard metrics as reported in Supplementary Figure 2. (F) Example single unit from left putamen showing raster plots with overlaid mean and SEM instantaneous spike rate traces during active dlPFC TMS (top) and sham dlPFC TMS (bottom). Gray shading around the vertical line at *t*=0 indicates the artifact period. Right panels display spike waveforms in color or gray with means in white corresponding to the spikes on the raster plots on the left, demonstrating consistent physiological appearing waveforms, without contamination by artifact, during active and sham dlPFC TMS. dlPFC, dorsolateral prefrontal cortex; EEG, electroencephalography; SEM, standard error of the mean; TMS, transcranial magnetic stimulation.

Given that TMS induces large electromagnetic artifacts in electrophysiological recordings that may obscure neural signals (Supplementary Figure 1), we adapted an artifact removal algorithm previously used for concurrent scalp EEG and TMS (Cline et al., 2021) for use with intracranial microwire recordings. This approach enabled detection of single-unit spiking in the post-stimulation period while preventing contamination of spike detection by the TMS artifact (Figure 1D; see Methods). After artifacts were removed from the raw broadband signal of each microwire channel, the data were bandpass filtered at 200-4000 Hz. Spikes were then detected offline using Higher Order Spectral Decomposition (Kovach et al., 2023; Kovach & Howard, 2019) and then clustered into single units. Single-unit quality and isolation was confirmed using established metrics (Supplementary Figure 2, Figure 1E). All spike waveforms were visually inspected to confirm physiological appearing morphology without sharp, large-amplitude transients characteristic of epileptiform activity. Across the 37 microwire bundles (8 recording wires each), at least one single unit was detected on 28 bundles. In total, we analyzed 185 single units across 296 microwires from all four participants. For each microwire channel, the artifactual period (onset -5 ms and mean and SD offset 12.7±14.9 ms relative to the TMS pulse) was excluded from analysis. Following artifact removal, single-unit spikes could be resolved as early as ∼8 ms after stimulation (Figure 1F, left). Spike waveforms recorded during active and sham TMS exhibited consistent morphology, confirming stable unit isolation without contamination by TMS artifact or epileptiform activity (Figure 1F, right).

### TMS modulates single-unit firing rates in widely distributed areas

To assess whether single-pulse TMS to the left dlPFC modulates single-unit activity, we compared post-stimulation firing rates to the pre-stimulation baseline separately in active and sham TMS conditions. Across the 185 single units, 50 units (27%) showed significantly increased firing rates (i.e., facilitation in spiking) within 1000 ms following single-pulse TMS (Figure 2A-B; *p_FDR_*<0.05, one-sided permutation test). We also found that 38/185 units (21%) exhibited significantly decreased firing rates (i.e., suppression in spiking) following stimulation. Only 2/185 (1%) of units exhibited both a significant increase and decrease. In contrast, in response to sham stimulation, 0/185 units exhibited a significant increase and 22/185 units exhibited a significant decrease. Of the 22 sham-suppressed units, 12 localized to right posteromedial Heschl’s gyrus (25 total units recorded in this region), suggesting that sham-related suppression may largely reflect auditory responses to the TMS click rather than direct TMS neuromodulatory effects (Supplementary Figure 3). Among Heschl’s gyrus single-units, active TMS suppressed 18/25. Collectively, there was a higher proportion of significantly modulated firing rates across all units in response to active compared to sham stimulation (*p_increase_*=0.0001 and *p_decrease_*=0.014, one-sided permutation test).

**Figure 2:**
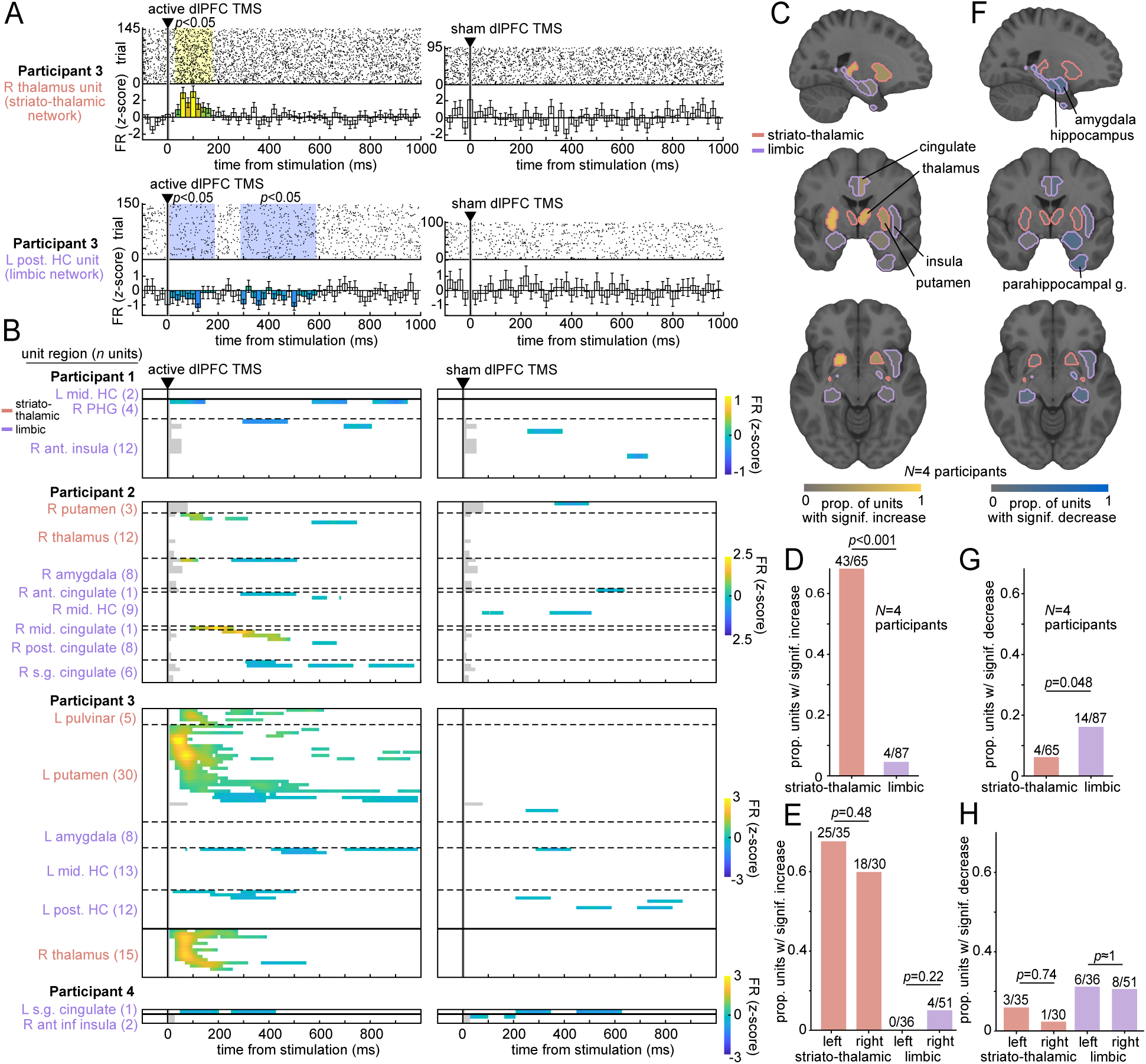
TMS facilitates neuronal spiking bilaterally in striato-thalamic regions and suppresses spiking bilaterally in limbic regions. (A) Example single unit rasters and *z*-scored firing rates (20 ms bin size) from two example single units showing contrasting responses to dlPFC TMS. Top shows a right thalamic single unit with significant facilitation of spike rates following active but not sham TMS. Bottom shows a left middle hippocampal single unit with significant suppression of spike rates following active but not sham TMS. Yellow and blue shading highlights significantly increased and decreased bins, respectively. Statistics were performed with random permutation tests with FDR correction for multiple comparisons across bins. (B) Heatmaps showing *z*-scored firing rates across all striato-thalamic and limbic single units from all participants (*n*=152 units, *N*=4 participants) during active dlPFC TMS (left) and sham dlPFC TMS (right). Each row represents one unit, with only statistically significant time bins displayed in color. Regions are separated by dashed lines and color-coded with striato-thalamic regions in red and limbic regions in purple. Within each region, units are sorted by the latency of their first significant firing rate change. Numbers in parentheses indicate the total units recorded in each region from each participant. Hemispheres are separated by solid horizontal black lines. Shaded gray denotes the artifactual period excluded from analysis. (C) Proportion of single units with significant firing rate increases following dlPFC TMS within each brain region, displayed on sagittal, coronal, and axial slices of the MNI152 standard brain template. Striato-thalamic regions in red and limbic in purple. (D) Quantification of the data in C reveals that a significantly higher proportion of striato-thalamic units (*n*=43/65) exhibited increased spike rates compared to limbic units (*n*=4/87), with *p*<0.001. Statistics were computed using a mixed-effects model with the participant as a random effect. (E) There was no significant difference in facilitation in striato-thalamic or limbic between hemispheres (*p*=0.48 and *p*=0.22, respectively). (F-H) Same as C-E except the proportion of single units with significant firing rate decreases following dlPFC TMS. Limbic regions showed a significantly greater suppression in spike rates compared to striato-thalamic (*n*=14/87 vs. *n*=4/65, respectively; *p*=0.048). There was no significant difference in suppression in striato-thalamic or limbic single units between hemispheres (*p*=0.74 and *p*≈1.0, respectively). FDR, false discovery rate; FR, firing rate; HC, hippocampus; PHG, parahippocampal gyrus.

### Single-unit facilitation localizes to bilateral striato-thalamic and suppression to bilateral limbic regions

We next assessed whether single-unit modulation during single TMS pulses is anatomically organized, given competing hypotheses that dlPFC stimulation may preferentially engage cortico-limbic affective pathways or striato-thalamic executive control networks. We classified each single unit by its anatomical location in striato-thalamic (putamen, thalamus, etc.), limbic (hippocampus, amygdala, subgenual cingulate, etc.), or other cortical and subcortical areas (Figure 2B, Supplementary Table 2). We found that TMS facilitated 43/65 (66%) of striato-thalamic single units compared to only 4/87 (5%) of limbic single units (Figure 2C-D; *p*<0.001, one-sided permutation test). TMS facilitated 25/35 left-hemispheric striato-thalamic single units and 18/30 right-hemispheric striato-thalamic single units, with no significant difference between these proportions (Figure 2E; *p*=0.48). By contrast, TMS facilitated 0/36 left-hemispheric limbic single units and 4/51 right-hemispheric limbic single-units, again with no significant hemispheric difference (*p*=0.22). Overall, TMS facilitation of single units was primarily in bilateral striato-thalamic regions, with no significant differences between ipsilateral (left) and contralateral (right) striato-thalamic or limbic single units.

Next, we examined the anatomical distribution of suppressed single units during TMS. We found that TMS suppressed the firing rate of 14/87 (16%) limbic neurons compared to 4/65 (6%) of striato-thalamic single units (Figure 2F-G; *p*=0.048; one-sided permutation test). Of these, TMS significantly suppressed 3/35 left hemispheric striato-thalamic single units and 1/30 right-hemispheric striato-thalamic single units, with no significant difference in these proportions (Figure 2H; *p*=0.74). Significant suppression was observed in 6/36 left-hemispheric limbic single units and in 8/51 right-hemispheric limbic single units, with no significant difference in the proportion of facilitated units between hemispheres (*p≈*1.0). Thus, in addition to facilitation, suppression was also observed bilaterally, with no significant differences between ipsilateral (left) and contralateral (right) striato-thalamic or limbic single units. Together, these results demonstrate that single-pulse dlPFC TMS produces anatomically dissociable effects: facilitation of striato-thalamic single units and suppression of limbic single units.

### Striato-thalamic single-unit facilitation precedes limbic suppression

To characterize the temporal dynamics of single-unit responses to dlPFC TMS, we computed the time course of modulated single units across the population (Figure 3A). TMS facilitated single-unit firing rates as early as ∼8 ms and peaked at ∼80-100 ms post-stimulation. Approximately 60% of striato-thalamic single units showed significant facilitation at this peak (Figure 3B). This facilitation then gradually decayed, returning to pre-stimulation baseline by ∼600 ms. Suppressed single units exhibited later and longer-lasting responses, peaking at ∼300 ms (Figure 3A,C), and returning to baseline by ∼700 ms. These results establish a temporal hierarchy in which striato-thalamic facilitation precedes limbic suppression.

**Figure 3:**
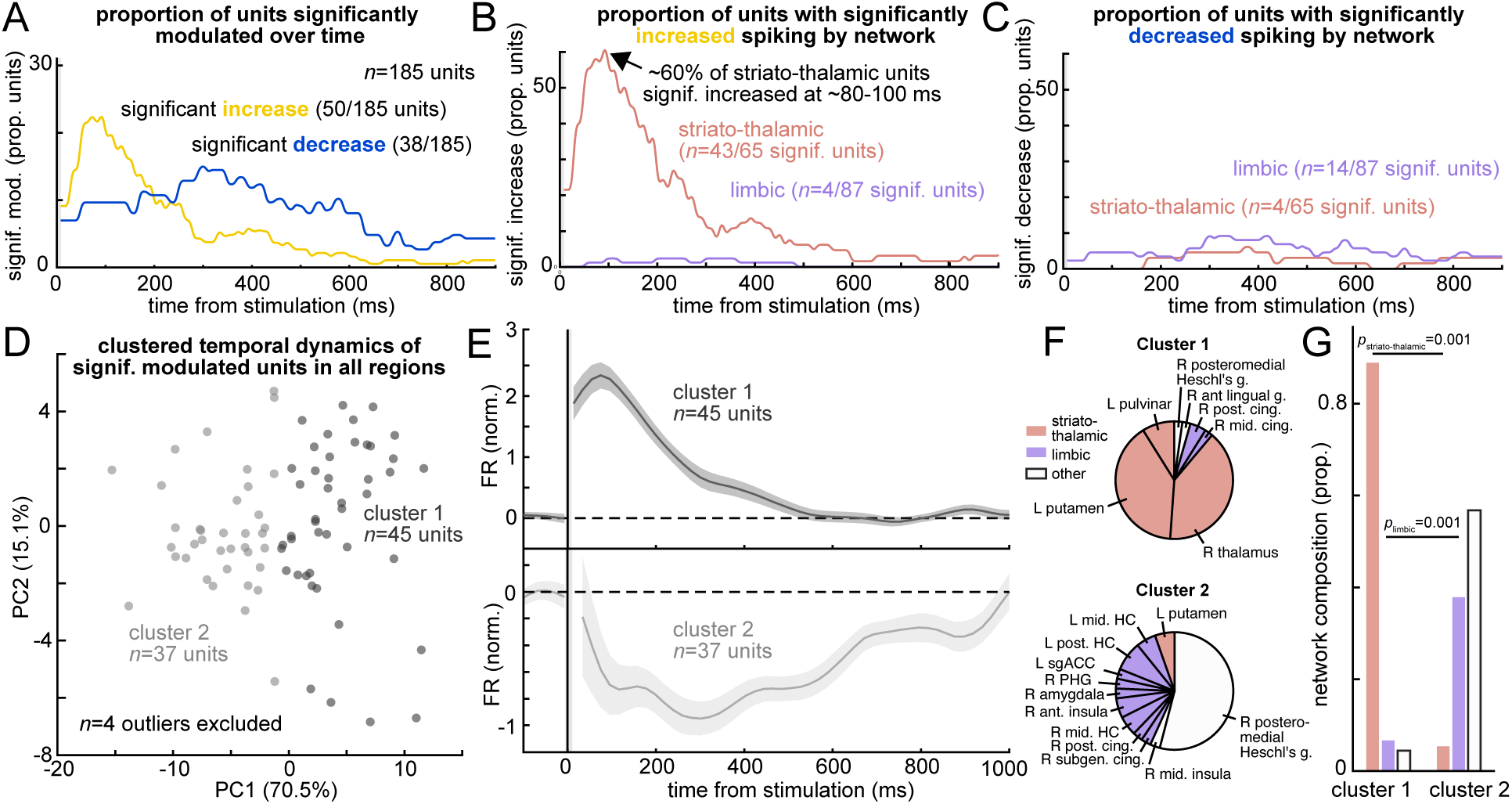
TMS facilitates early striato-thalamic single-unit spiking and suppresses late limbic single-unit spiking. (A) Time course showing the proportion of all single units (*n*=185) from all participants (*N*=4) with significant firing rate modulations following active dlPFC TMS. Single units with significant increases are shown in yellow and those with significant decreases are shown in blue. (B) Time course showing the proportion of striato-thalamic single units (*n*=65; red) and limbic single units (*n*=87 units; purple) with significant firing rate increases following active dlPFC TMS. (C) Same as B except proportions with significant firing rate decreases. (D) Principal component analysis of normalized firing rate responses across all significantly modulated single units (*n*=82 units, 4 outliers excluded). Clustering using *k*-means identified two distinct temporal response patterns, with cluster 1 shown in dark gray (*n*=45 units) and cluster 2 shown in light gray (*n*=37 units). (E) Mean and SEM normalized firing rates for each cluster in D. Top shows cluster 1, which exhibits facilitation peaking at ∼80 ms and decaying until ∼600 ms. Bottom shows cluster 2, which exhibits suppression peaking at ∼300 ms and sustained until ∼1000 ms. (F) Pie charts show the network composition of the two clusters by unit region. (G) Quantification of network (striato-thalamic, limbic, or other) composition by clusters in D-E reveals that cluster 1 with early peaking facilitation contains significantly more striato-thalamic single units (*p*=0.001) whereas cluster 2 with late suppression contains significantly more limbic single units (*p*=0.001). PC, principal component.

### Single units exhibit two response phenotypes: rapid facilitation and slower suppression

To characterize response dynamics, we performed unsupervised clustering of single-unit temporal dynamics. We included all single units with significant modulation in either direction (*n*=86 units) and computed normalized firing rates in 20 ms bins from 0-1000 ms post-stimulation. Principal component analysis (PCA) reduced these temporal profiles to three dimensions, capturing 92.7% of variance. Four outlier units were removed using an Isolation Forest algorithm on the PCA scores. Clustering was then performed using *k*-means, where the optimal number of clusters (*k*) was determined through a consensus approach integrating four validation indices (see Methods). The clustering was run 50 times to ensure a stable solution, which identified two distinct response phenotypes (Figure 3D). One cluster (*n*=45 units) exhibited rapid facilitation with peak firing at ∼80 ms, followed by gradual decay back toward baseline by ∼600 ms (Figure 3E, top). The second cluster (*n*=37 units) exhibited a later suppression, which peaked ∼300 ms and was sustained until ∼1000 ms post-stimulation (Figure 3E, bottom). The first cluster demonstrating facilitation included 40/42 striato-thalamic (including thalamus and putamen) and 3/17 limbic units (Figure 3F). The second cluster demonstrating suppression included 14/17 limbic (including hippocampus, amygdala, anterior insula, and subgenual cingulate) and 2/42 striato-thalamic units. There were significantly more striato-thalamic units in the facilitation than suppression cluster (Figure 3G; *p*=0.001, generalized linear-mixed effects model), and by contrast significantly more limbic units in the suppression than facilitation cluster (*p*=0.001). Performing dimensionality reduction using *t*-distributed Stochastic Neighbor Embedding instead of PCA also yielded two response phenotypes, one demonstrating facilitation and the other suppression in single-unit spiking in response to TMS (Supplementary Figure 4A-C). Performing clustering using a Gaussian mixture model instead of *k*-means also yielded these two phenotypes (Supplementary Figure 4D-F). Together, these complementary approaches confirm the robustness of the identified facilitation and suppression response phenotypes across multiple analytical methods.

### TMS facilitates striato-thalamic putative interneurons and suppresses widespread putative pyramidal cells

Next, we investigated whether TMS differentially modulates excitatory and inhibitory neurons. Single units were classified as putative pyramidal cells (PY, *n*=102) or interneurons (IN, *n*=83) according to their spike waveform and timing features using Gaussian mixture modeling of normalized features: trough-to-peak duration, full width at half maximum, burst index, and firing rate across all four participants (Figure 4A-B; see Methods). PY exhibited significantly longer full-width at half-maximum, lower burst index, lower firing rate, and larger CV2 compared to putative interneurons (Figure 4C; all *p*<0.001, linear-mixed effects models). These metrics are consistent with those established in the single unit literature (Barthó et al., 2004; McCormick et al., 1985) including human studies with single units sampled from cortical and subcortical regions (Agopyan-Miu et al., 2023; Dickey et al., 2021; Kussovska et al., 2025; Peyrache et al., 2012). Single-pulse TMS had opposing effects on the two cell types, facilitating spiking in a significantly greater proportion of IN than PY (Figure 4D; *p*<0.001, one-sided permutation test) and suppressing spiking in a significantly greater proportion of PY than IN (*p*=0.016). These differential effects showed distinct anatomical patterns, with IN recruitment significantly more prevalent in striato-thalamic compared to limbic regions (Figure 4E; *p*<0.001), while PY suppression showed a non-significant trend toward greater prevalence in limbic compared to striato-thalamic regions (Figure 4F; *p*=0.077).

**Figure 4:**
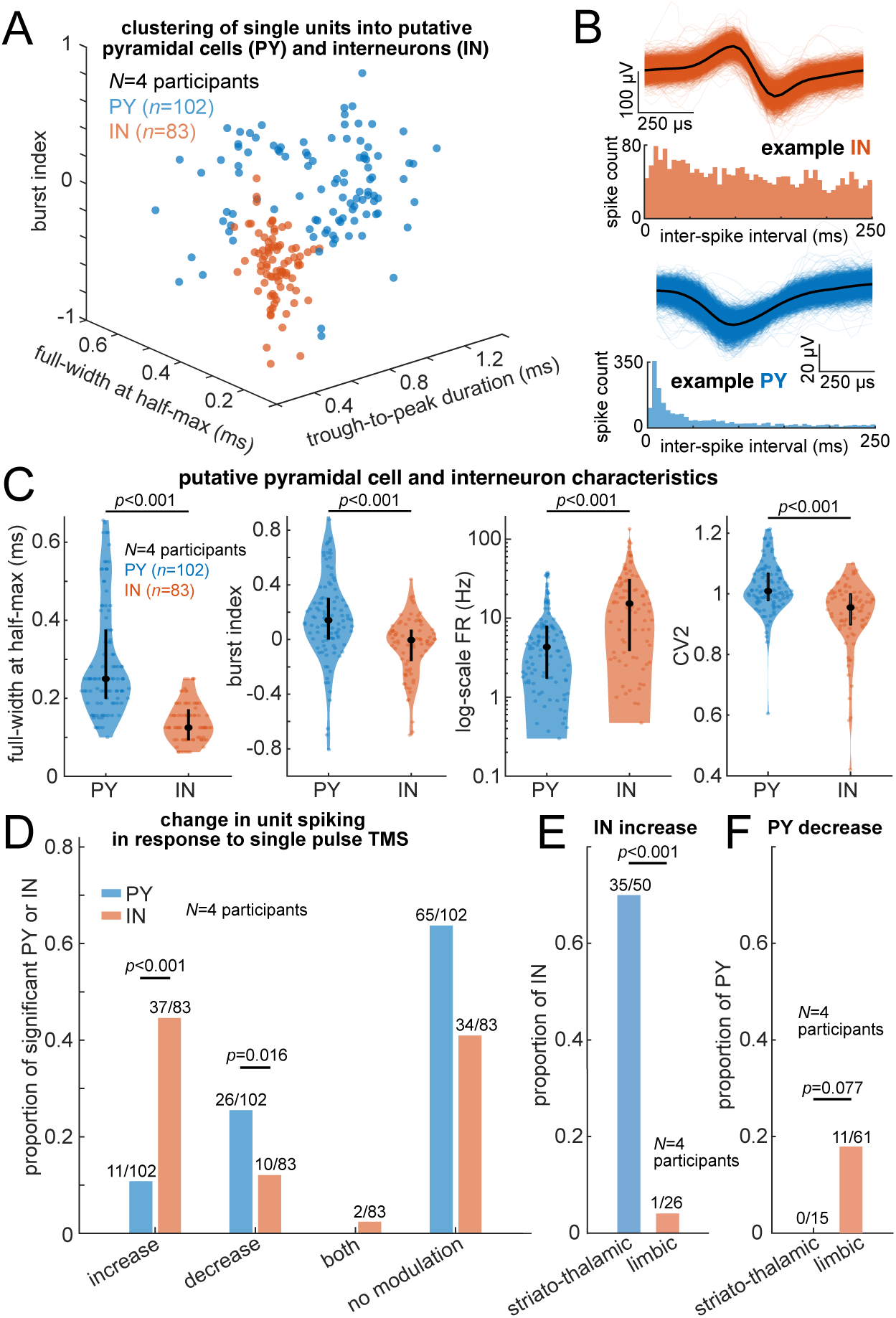
Single-pulse TMS recruits putative interneurons in striato-thalamic regions and suppresses putative pyramidal cells in widespread areas. (A) Single units clustered into putative pyramidal cells (PY, *n*=102) and interneurons (IN, *n*=83) using Gaussian mixture modeling of normalized waveform features: trough-to-peak duration, full width at half maximum, burst index, and firing rate. Three of four features shown. (B) Spike waveforms and inter-spike interval distributions from an example IN (top) and PY (bottom). Individual spikes in color and mean in black. (C) PY spike waveform averages have a significantly longer full-width at half-maximum, lower burst index, lower firing rate, and larger CV2 compared to IN (all *p*<0.001, one-sided permutation test). Black circles show median and vertical bars show interquartile range. (D) Single-pulse TMS increases spiking in a greater proportion of IN than PY and decreases spiking in a greater proportion of PY than IN (*p*<0.001 and *p*=0.016 respectively, permutation test). (E) IN recruitment is significantly more prevalent in striato-thalamic than limbic regions (*p*<0.001). (F) PY suppression shows a non-significant trend toward greater prevalence in limbic vs. striato-thalamic regions (*p*=0.077). CV2, coefficient of variation 2; IN, interneuron; PY, pyramidal cell.

### Single-unit responses are positively correlated with striato-thalamic and anti-correlated with limbic network activity

A central question remains as to whether single-unit modulations reflect local circuit dynamics or large-scale network engagement. To address this, we simultaneously recorded intracranial TMS-evoked potentials (iTEPs) from macroelectrode bipolar re-referenced channels (Figure 5A-B) in response to the same TMS single pulses used for single-unit analyses. iTEPs map intracranial network connectivity in response to TMS (Hassan, 2025; Solomon et al., 2024, 2026; Wang et al., 2024). Within the associative cortico-striato-thalamic network, single-unit recordings were obtained from the striatum and thalamus, whereas macroelectrodes also captured field potentials from associative cortical regions (Supplementary Table 2). The primary analyses of correlations between single-unit modulations and iTEPs (Figure 5) required adequate simultaneous sampling of both iTEPs across cortico-striato-thalamic and limbic areas and single units across striato-thalamic and limbic areas (see Methods). Participant 3 met these criteria and was included for the primary analyses. Of 61 bipolar re-referenced macroelectrode channels, 25 were classified as cortico-striato-thalamic (Figure 5A), 6 as limbic, and 30 as other. iTEP amplitudes (*z*-scored root mean squared, 50-500 ms post-stimulation; see Methods) did not significantly differ between cortico-striato-thalamic and limbic networks (Figure 5C; *p*=0.10, *z*=1.6, two-sided Wilcoxon rank-sum test). Among 83 single units from these regions, 50 were striato-thalamic (42 responsive and 8 non-responsive to active TMS) and 33 were limbic (6 responsive and 27 non-responsive); no single units fell outside these regions (Figure 5D, inset). Striato-thalamic single units with a significant spike rate modulation (either direction) had greater spike rate modulation (Poisson *z*-score, 50-500 ms) than limbic single units with a significant spike rate modulation (Figure 5D; *p*<0.001, *z*=3.6, two-sided Wilcoxon rank-sum test).

**Figure 5:**
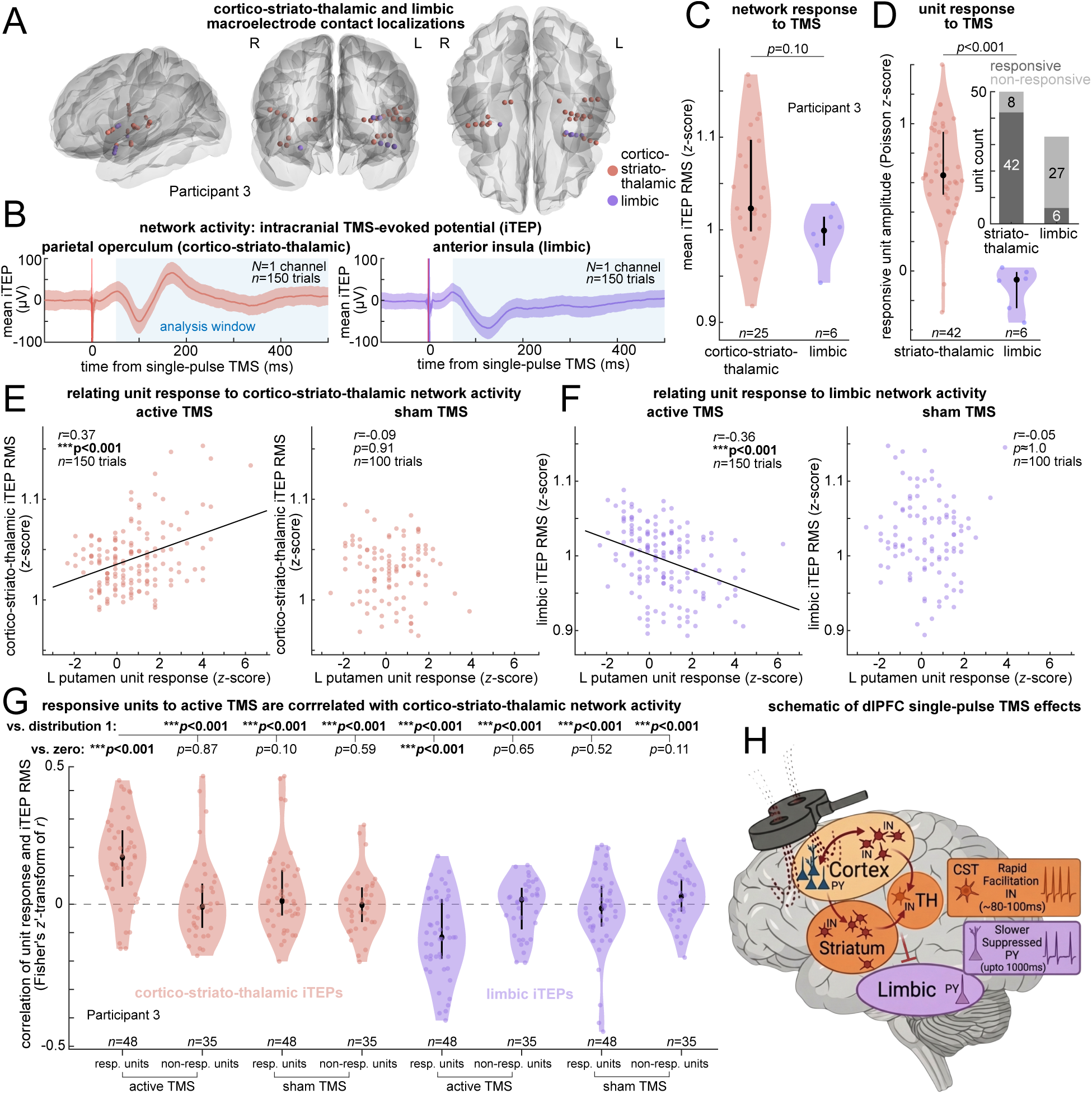
Single-unit responses are correlated with striato-thalamic and anti-correlated with limbic network recruitment. (A) Macroelectrode contact locations (participant 3) on fsaverage template brains, color-coded by network affiliation with cortico-striato-thalamic in red and limbic in purple. (B) Example mean and SEM iTEP traces from an example macroelectrode bipolar re-referenced channel in the parietal operculum within the cortico-striato-thalamic network (left, *n*=150 trials of single-pulse TMS to dlPFC) and anterior insula within the limbic network (right, *n*=150 trials). Shaded box indicates the 50 to 500 ms period used for the subsequent analyses. (C) iTEP RMS amplitude from 50 to 500 ms in cortico-striato-thalamic (*n*=25 macroelectrode bipolar re-referenced channels, red) and limbic (*n*=6 channels, purple) networks (*p*=0.10; Wilcoxon rank-sum test). Black circles show median and black vertical bars show interquartile range. (D) Single-unit response magnitudes (Poisson *z*-score) compared to baseline between striato-thalamic (*n*=42 units) and limbic (*n*=6 units) networks (*p*=0.003). Inset shows the number of responsive and non-responsive units in striato-thalamic vs limbic regions. (E) Scatter plots showing the relationship between cortico-striato-thalamic iTEP RMS and single-unit response magnitude for an example left putamen unit within the striato-thalamic network during active TMS (left, *r*=0.37, *p*_FDR_<0.001, *n*=150 trials) and sham TMS (right, *r*=-0.09, *p*_FDR_=0.91, *n*=100 trials). (F) Same as D but showing the relationship between limbic network iTEP RMS and the same left putamen unit response magnitude during active TMS (left, *r*=-0.36, *p*_FDR_<0.001, *n*=150 trials) and sham TMS (right, *r*=-0.05, *p*_FDR_≈1.0, *n*=100 trials). (G) Correlations between single-unit responses and iTEP RMS across different conditions. Data are grouped by responsive units (resp. units), non-responsive units (non-resp. units) for the cortico-striato-thalamic iTEP network (red, left four violins) and limbic iTEP network (purple, right four violins) during active and sham dlPFC TMS. Critically, only responsive units during active TMS exhibited robust positive correlations with cortico-striato-thalamic iTEPs (leftmost distribution, *n*=48 units), significantly exceeding all other conditions (all *p*_FDR_<0.001, one-sided Wilcoxon rank-sum test). (H) Schematic illustrating the mechanistic framework revealed by simultaneous single-unit and network recordings. Single-pulse TMS to the left dlPFC preferentially recruits the striato-thalamic network, evidenced by facilitation of single-unit spiking in striatum and thalamus that is tightly coupled to cortico-striato-thalamic network activity on a trial-by-trial basis. This selective engagement of executive control pathways occurs preceding a suppression of limbic single-unit activity. These findings suggest that dlPFC TMS may rebalance large-scale control networks by strengthening striato-thalamic gating mechanisms that regulate top-down inhibitory control over limbic reactivity, rather than through direct normalization of limbic hyperactivity. CST, cortico-striato-thalamic network; iTEP, intracranial TMS-evoked potential; RMS, root mean square; TH, thalamus. ***p<0.001.

To test whether single-unit firing couples to network activity, we computed trial-by-trial correlations between iTEP amplitude and single-unit firing rate modulation (see Methods). In an example striato-thalamic single unit in the left putamen, the firing rate modulation was significantly positively correlated with cortico-striato-thalamic iTEP amplitude following active TMS (Figure 5E, left; *r*=0.37, *p*_FDR_<0.001) but not sham TMS (Figure 5E, right; *r*=-0.09, *p*_FDR_=0.91). The same unit was significantly anti-correlated with limbic iTEP during active (Figure 5F, left; *r*=-0.36, *p*_FDR_<0.001) but not sham TMS (Figure 5F, right; *r*=-0.05, *p*_FDR_≈1.0). Across all regions, responsive units (*n*=48 responsive; *n*=35 non-responsive; comprising both striato-thalamic and limbic units) during active TMS were significantly correlated with cortico-striato-thalamic network response (Figure 5G; mean *r_z_*=0.15±0.15, *p*_FDR_<0.001, *z*=5.0, two-sided Wilcoxon signed-rank test). Conversely, responsive single units during active TMS were significantly anti-correlated with limbic iTEPs (mean *r_z_*=-0.10±0.14, *p*_FDR_<0.001, *z*=4.3). There were no significant correlations involving non-responsive units or sham TMS (*p*_FDR_=0.10-0.87, *z*=0.5-1.6). Furthermore, responsive single units during active TMS exhibited significantly greater positive correlations with cortico-striato-thalamic iTEPs than all other conditions: non-responsive single units during active TMS (*p*_FDR_<0.001, *z*=4.0, one-sided Wilcoxon rank-sum test), responsive single units during sham TMS (*p*_FDR_<0.001, *z*=3.9), non-responsive single units during sham TMS (*p*_FDR_<0.001, *z*=4.7), and all comparisons with limbic iTEPs (responsive active: *p*_FDR_<0.001, *z*=6.7; non-responsive active: *p*_FDR_<0.001, *z*=5.2; responsive sham: *p*_FDR_<0.001, *z*=5.1; non-responsive sham: *p*_FDR_<0.001, *z*=4.1). These correlations did not differ significantly between PY and IN in any condition (Supplementary Figure 5; all *p*_FDR_≥0.53). Substituting iTEP amplitude with TMS-evoked high gamma (70-190 Hz) amplitude yielded no significant correlations with responsive single-unit modulations during active TMS (Supplementary Figure 6; all *p_FDR_*≥0.05), indicating that the observed single-unit coupling was specific to lower-frequency network activity captured by the iTEP. Together, these findings demonstrate that trial-by-trial responsive single-unit modulations in striato-thalamic and limbic regions to active TMS are positively correlated with striato-thalamic network activity and anti-correlated with limbic network activity.

### Striato-thalamic single units drive network-specific correlations

To examine whether the correlations between responsive single-unit modulations and network-level iTEPs were specific to striato-thalamic single units, we computed trial-by-trial correlations separately for striato-thalamic and limbic single units against each network’s (cortico-striato-thalamic and limbic) iTEP amplitude (Supplementary Figure 7). In participant 3, responsive striato-thalamic single units during active TMS exhibited a significant positive correlation with cortico-striato-thalamic iTEPs (mean *r_z_*=0.18±0.15, *p*_FDR_<0.001, two-sided Wilcoxon signed-rank test) and a significant anti-correlation with limbic network iTEPs (mean *r_z_*=-0.12±0.14, *p_FDR_*<0.001), whereas there were no significant correlations with either cortico-striato-thalamic or limbic iTEPs for limbic units, non-responsive units, or sham TMS (Supplementary Figure 7A; all *p_FDR_*=0.13-0.98). This indicates that the correlations observed in Figure 5G were driven by striato-thalamic units which were responsive to active TMS. An exploratory analysis in participant 2, the only other participant with adequate macroelectrode coverage of both networks but with a limited number of responsive striato-thalamic single units (*n*=3) and responsive limbic single units (*n*=12), showed a non-significant positive correlation between responsive striato-thalamic single units and cortico-striato-thalamic iTEPs (Supplementary Figure 7B; mean *r_z_*=0.11±0.07, *p_FDR_*=0.49). In participant 2, there was no significant anti-correlation as was observed in participant 3 (mean *r_z_*=0.10±0.08, *p_FDR_*=0.49), though the smaller unit yield limits interpretation of this null finding. Together, these analyses demonstrate a dissociation in which responsive single-unit modulations are positively correlated with striato-thalamic but anti-correlated with limbic network activity on a trial-by-trial basis, driven by striato-thalamic single units, with partial though inconclusive support from a second participant.

## Discussion

### Summary of key findings: single-neuron responses to dlPFC TMS

This study provides the first investigation of single-neuron responses to dlPFC TMS in humans. By applying single pulses of TMS to the left dlPFC, we found that stimulation modulates single-neuron spiking in deep brain regions across both ipsilateral and contralateral hemispheres. Our artifact cleaning approach enabled resolution of neuronal spiking as early as ∼8 ms post-stimulation, a critical advance that has previously required extensive hardware modifications not yet feasible in humans (Mueller et al., 2014). The data reveal a fundamental dissociation in how stimulation engages distributed networks. TMS rapidly facilitated single-neuron spiking in striato-thalamic regions, with responses emerging as early as ∼8 ms, peaking at ∼80-100 ms, and lasting up to ∼600 ms. By contrast, TMS suppressed spiking in limbic neurons in a delayed and sustained manner, with effects lasting to ∼1000 ms. Cellular classification revealed that single pulse TMS recruits INs in striato-thalamic regions and inhibits PYs in widespread areas. Single-neuron spiking modulations were correlated with macroelectrode TMS-evoked responses in the striato-thalamic network and were anti-correlated with evoked responses in the limbic network. These findings are consistent with dlPFC TMS selectively recruiting executive control circuits and exerting inhibitory gating over limbic structures.

### Single-neuron responses to TMS in animal models

Our findings align with and extend those from the majority of single-neuron recordings performed during TMS in animal models, where single neurons have primarily been recorded at the local cortical area being stimulated. In *in vitro* mouse organotypic entorhino-hippocampal slice cultures, high-frequency (10 Hz) repetitive magnetic stimulation facilitated an increase in the glutamatergic synaptic strength of CA1 PYs and remodeling of dendritic spines (Vlachos et al., 2012). In cat visual cortex, short TMS pulse trains activated spontaneous local single-neuron spiking for ∼1 minute, with suppression of spiking during visual stimuli for 5-10 minutes, as well as decreased phase-locking of spontaneous spikes to multiple frequency bands including theta (Allen et al., 2007). In the anesthetized cat primary visual cortex, single-pulse TMS facilitated local single-neuron spiking ∼200-500 ms after stimulation followed by a slower suppression (Moliadze et al., 2003). In alert non-human primates, single-pulse TMS to the frontal eye field modulated local spiking of single putative axons, INs, and PYs, with increased activity lasting at least 100 ms (Mueller et al., 2014). In awake behaving macaque monkeys, single-pulse TMS to parietal cortex elicited a transient activation in single-neuron spiking within 2 mm of the stimulation site for 10-80 ms after stimulation, with some neurons showing subsequent suppression for 50 ms, followed by a second phase of excitation lasting up to 250 ms (Romero et al., 2019). Across these studies there was a median of 48 single-neurons recorded (range=18-476) in response to TMS. In summary, animal studies reveal that TMS typically triggers an early facilitation followed by suppression in neurons near the stimulation site, with precise response patterns varying across experimental conditions. Our findings in humans align with this fundamental principle, demonstrating a rapid facilitation in deep brain neuronal firing (Figures 1F, 2A-B, 3A,B,F). In addition to recording from distributed regions distant from the stimulation site, we extend the work in animal models by revealing network- and cell-type-specific response profiles that have not been previously described.

### Rapid bilateral engagement of striato-thalamic pathways

Consistent with recent TMS-iEEG work demonstrating that single-pulse dlPFC TMS provokes widespread oscillatory changes in fronto-limbic networks including the hippocampus and anterior cingulate cortex (Solomon et al., 2024, 2026), our single-neuron recordings reveal the cellular substrates underlying these distributed network effects. The measurement of neuronal facilitation as early as ∼8 ms post-stimulation in regions distant from the dlPFC suggests that TMS engages fast-conducting mono- or oligo-synaptic pathways. This rapid facilitation was prominent in the putamen and thalamus, nodes of the striato-thalamic loop which are critical for executive function (Alexander et al., 1986). These findings are consistent with fMRI evidence that dlPFC-striatal functional connectivity predicts TMS treatment response (Avissar et al., 2017), and suggest that rapid neuronal facilitation within striato-thalamic circuits may underlie this relationship. Strikingly, we observed parallel activation in the right thalamus, revealing that unilateral prefrontal stimulation is capable of rapid interhemispheric propagation facilitating neuronal modulation in bilateral striato-thalamic pathways. Given that depression is characterized by network-wide dysregulation rather than focal pathology (Liston et al., 2014), this distributed pattern of engagement is relevant for understanding how prefrontal stimulation influences mood-related circuitry. Moreover, early facilitation in striato-thalamic neurons was correlated with network activity across the cortico-striato-thalamic network, demonstrating that TMS drives coordinated activity across distributed executive control circuits. More broadly, the unexpectedly high proportion of responsive neurons we observed across both hemispheres suggests that dlPFC TMS engages widespread neural populations well beyond the stimulation site, potentially explaining why varied targeting approaches (e.g., 5.5 cm rule, Beam F3, and subgenual anterior cingulate anti-correlated sites) yield comparable clinical outcomes in the treatment of depression (Terao & Kodama, 2025; Trapp et al., 2023).

### Recruitment of striato-thalamic putative interneurons and suppression of widespread putative pyramidal cells

Single-pulse TMS to dlPFC drives activation of striato-thalamic INs while subsequently suppressing PYs, which is consistent with the literature on cortico-striato-thalamic neurophysiology. Recent evidence suggests that cortical stimulation engages cortico-thalamo-cortical loops through GABAergic IN-mediated thalamic hyperpolarization and rebound (Russo et al., 2025), driving each cortico-thalamic node at its natural frequency (Rosanova et al., 2009). This IN recruitment may enable rapid and coordinated modulation of network activity, potentially explaining the speed and bilateral extent of the observed striato-thalamic responses. This aligns with recent rodent work where accelerated intermittent theta burst stimulation (iTBS) facilitated plasticity in cortico-striatal pathways (Gongwer et al., 2025), suggesting the single-pulse recruitment we observed may include areas where plastic changes are induced by repetitive stimulation.

### Dissociable mechanisms of striato-thalamic facilitation and limbic suppression suggest top-down inhibitory gating

While striato-thalamic single neurons showed rapid facilitation positively coupled to cortico-striato-thalamic network activity and inversely related to limbic network activity, limbic neurons exhibited delayed and sustained suppression without significant coupling to either network. This dissociation, observed most robustly in the participant with the most extensive sampling, suggests distinct mechanisms: striato-thalamic neuronal facilitation may result from direct recruitment of cortico-striato-thalamic pathways, whereas limbic neuronal suppression may arise indirectly through striato-thalamic circuit engagement rather than direct cortico-limbic propagation. Collectively, these data provide a mechanistic explanation for prior observations that dlPFC stimulation sites effective for clinical improvement are anti-correlated with the subgenual cingulate (Fox et al., 2012). Rather than direct excitatory propagation to limbic regions, our findings support a top-down control model where dlPFC stimulation drives cortico-striato-thalamic loops to gate limbic reactivity. The recruitment of striato-thalamic INs may serve as the cellular trigger for this remote modulation. Our finding that trial-by-trial striato-thalamic facilitation is inversely related to limbic network activity provides a potential single-neuron substrate for the network-level anti-correlation observed by Fox and colleagues, potentially occurring via IN-mediated gating of limbic activity. Our data suggest that left dlPFC stimulation first facilitates striato-thalamic INs, which may subsequently inhibit limbic neurons, as evidenced by the temporal precedence of striato-thalamic facilitation over limbic suppression. Our results therefore support the hypothesis that TMS may re-balance activity between executive control and limbic networks. Of note, responses to single pulses of cortical stimulation have been shown to predict the spatial distribution of effects induced by repetitive stimulation (Huang et al., 2019, 2024; Keller et al., 2018), suggesting that the single-pulse TMS-evoked neuronal responses observed here may reflect recruitment patterns evoked during therapeutic repetitive TMS protocols.

### Limitations and future directions

In summary, we provide the first direct evidence at the single-neuron level for the long-held hypothesis that TMS selectively engages and modulates large-scale brain circuits in humans. Our results suggest that single-pulse TMS recruits striato-thalamic INs followed by suppression of limbic neuronal activity and widespread gating of PY. While this study provides the first view into the single-neuron effects of dlPFC TMS in humans, several limitations should be noted. Neuronal sampling was sparse and differed between participants, which is inherent to these opportunistic recordings. As data were collected from patients with epilepsy, the possibility of pathological network remodeling cannot be fully excluded. However, the sampling of non-epileptogenic areas, absence of epileptiform activity in the data included in the study, along with the consistency of our findings across patients with differing epileptic foci suggests a generalizable physiological mechanism. In addition, this study assessed neuronal activity by measuring spike rate modulations, which does not consider that other types of modulation, for example phase-locking of neuronal spiking to rhythms in the LFP, may be relevant to these circuits. Despite these limitations, these exceptionally rare recordings have offered an unprecedented window into the core mechanisms of TMS. Having established how single-pulse TMS modulates neuronal activity, a critical future direction will be to determine how therapeutic protocols like iTBS affect neuronal activity and plasticity within these networks.

## Methods

### Participant selection

Participants with intractable epilepsy undergoing clinical monitoring for seizure focus localization were included in this study if they received dlPFC TMS and were implanted with recorded microwires that had single units isolated offline (Supplementary Table 1). Depression status was not an inclusion or exclusion criterion. Among nine participants receiving TMS while implanted with microwires, four were included in the study. One participant was excluded due to pathological electrophysiological activity in the majority of recording sites based on visual inspection. The remaining four were excluded due to either amplifier saturation affecting all high impedance contacts or significant noise contaminating the recordings such that single units were not able to be resolved. In certain cases, this included noise caused by the presence of a separate active recording system in the room. All participants in this study gave fully informed written consent for their data to be used for research purposes. This study was approved by the local Institutional Review Board at the University of Iowa in accordance with the principles of the Declaration of Helsinki.

### Electrophysiological recordings

Stereo-electroencephalography was recorded using depth electrodes (Ad-Tech Medical, Oak Creek, WI) which were implanted according to clinical indications for localizing epileptogenic zones. Intracranial recordings during TMS were conducted following established protocols (Hassan, 2025; Wang et al., 2024). Macroelectrode recordings were amplified, filtered (ATLAS, Neuralynx, Bozeman MT; 0.7-800 Hz bandpass), and digitized at a sampling rate of 8 kHz (participant 1), 16 kHz (participant 2), 32 kHz (participant 3), or 40 kHz (participant 4). These recordings were downsampled offline to 1000 Hz with anti-aliasing and bipolar referenced using adjacent contacts on the same probe. Macroelectrode recording contacts were excluded from analysis if stimulation artifacts saturated the amplifier, if electrodes were placed within an identified seizure onset zone, if the bipolar re-referenced channel did not localize to gray matter, or if recordings were contaminated by non-neural noise indicative of poor connection or placement outside the brain.

Microwire recordings were collected simultaneously during macroelectrode recordings. Of the 58 depth electrode shafts implanted across four patients, 37 included nine high impedance platinum-iridium microwires of 39 µm diameter, eight of which were insulated (Figure 1B). The microwires extended beyond the distal end of the macroelectrode shaft at lengths ranging from 2-4 mm, with specific lengths determined by the distance between the terminal macro-contact and the intended recording site. Prior to insertion, each microwire was manually separated into a splayed configuration in the surgical suite. Neurophysiological signals were acquired using a Neuralynx Atlas System (Neuralynx, Bozeman, MT). The eight recording microwires were referenced to the ninth uninsulated wire. Signals from the high impedance microwires were amplified via a head-mounted preamplifier unit (ATLAS-HS-36-CHET-A9, Neuralynx, Bozeman, MT) before digital conversion. Microwire data were sampled at 32 kHz for participants 1-3 and 40 kHz for participant 4, with bandpass filtering from 0.1 to 8000 Hz. Single-unit data and macroelectrode recordings were subsequently analyzed in Matlab 2024b (MathWorks, Natick, MA) and Python 3.12.2 (Python Software Foundation, Wilmington, DE).

### Anatomical localization

Intracranial electrodes were localized following previously described procedures (Wang et al., 2024). High-resolution T1-weighted MRI scans were acquired for all participants at two timepoints: within two weeks prior to electrode implantation and following the surgical implantation of electrodes. Imaging was performed on a 3T Siemens TIM Trio system equipped with a 12-channel head coil. The MPRAGE sequence parameters included in-plane resolution of 0.78 × 0.78 mm with 1.0 mm slice thickness, TR of 2.53 s, and TE of 3.52 ms. Anatomical localization of recording sites was determined by registering pre-operative and post-operative structural scans using 3D linear registration (Functional MRI of the Brain Linear Image Registration Tool; (Jenkinson et al., 2012)) combined with custom Matlab code. Individual electrode coordinates were transformed into MNI space through normalization to a standard template for visualization purposes.

FreeSurfer (Fischl, 2012) was used to map electrode locations onto a standardized set of coordinates across participants, which were then labeled according to their location within the Desikan-Killiany-Tourville anatomical atlas (Desikan et al., 2006). Anatomical localization of the microwire bundles was confirmed by visual inspection. Visualization of microwire bundle and macroelectrode channels on 3D FreeSurfer fsaverage template brains and MNI152 brain slices was performed in Python.

Regarding network assignments (i.e., cortico-striato-thalamic, striato-thalamic, and limbic), we adopted an anatomical classification approach, assigning recording sites to networks based on their position within these established circuit architectures. Our single-unit analyses (Supplementary Table 1) focused on subcortical nodes of the executive control circuit (i.e., the striatum and thalamus), while the macroelectrode network analyses presented in Figure 5 additionally incorporated cortical recording sites (Supplementary Table 2). The term ‘striato-thalamic’ is used throughout to refer collectively to the subcortical components of this prefrontal-associated loop, consistent with prior nomenclature distinguishing executive control circuits from limbic circuits centered on orbitofrontal and anterior cingulate cortices (Alexander et al., 1990).

### Transcranial magnetic stimulation

For stimulation, we used a MagVenture X100 system with a figure-of-eight liquid-cooled Cool-B65 A/P coil (MagVenture, Alpharetta, GA). Stimulation pulses were biphasic sinusoidals with a pulse width of 290 microseconds, with stimulator output set at a percentage of each participant’s motor threshold (Supplementary Table 1). Pulses were delivered at 0.33-0.4 Hz, corresponding to 2.5-3 s inter-stimulation intervals using automated scripting from the BEST toolbox (Hassan et al., 2022). TMS experiments were conducted 12-19 days after implantation, once anti-seizure medication had been re-initiated. Due to the microwire recordings being obtained this long after implantation, a reduced single-unit yield is expected. Neuronavigation using frameless stereotaxy was guided with Brainsight v2.4.7 software (Rogue Research, Inc., Montreal, Quebec, Canada) supplied with the pre-implantation T1/MPRAGE anatomical scan. TMS was targeted to the left dlPFC using a coordinate aligned with the modified Beam F3 targeting method (Beam et al., 2009; Trapp et al., 2020). Stimulation parameters were recorded in Brainsight during all experimental trials. Motor thresholds were determined starting with the hand knob of the motor cortex as a target, beginning at 50% machine output and adjusted in small increments until movements were observed in 50% of trials. As specified in Supplementary Table 1 control conditions involved scalp electrical stimulation to simulate the somatosensory effects of TMS (in one sham block for participant 3) and/or involved auditory masking using TMS Adaptable Auditory Control (Russo et al., 2022; Trapp et al., 2024).

### Artifact cleaning

While hardware methods utilized in animal models exist to resolve single-unit spikes within several milliseconds of the TMS pulse, these approaches are not currently feasible in humans (Mueller et al., 2014; Tischler et al., 2011). To address stimulation artifacts, we implemented a comprehensive multi-stage artifact detection and correction protocol. We first detected the bounds of stimulation artifacts and any high-frequency, large-amplitude noise around stimulation times on a per-trial basis (Supplementary Figure 1). For each stimulus pulse, an initial artifact window was defined by identifying periods of signal saturation (at least 10 consecutive identical sample values), with a minimum enforced window of 5 ms before and after the pulse. This window was then extended by identifying residual high-frequency artifacts through bandpass filtering (800-1200 Hz), calculating the signal envelope via Hilbert transform, and *z*-scoring relative to a pre-stimulus baseline (-50 to -5 ms). When the post-stimulus *z*-score exceeded a threshold of 10, the artifact window was extended to the point where the *z*-score returned to 0. Data within this final combined artifact window were replaced using autoregressive extrapolation with 10 ms pre- and post-fit durations, following established interpolation methods for simultaneous TMS and scalp EEG (Cline et al., 2021). The cleaned broadband signal was then used for subsequent spike detection and sorting, employing Higher Order Spectral Decomposition (see below for details) to distinguish physiological action potentials from potential remaining noise or artifact (Berger et al., 2026; Kovach et al., 2023; Kovach & Howard, 2019). Our approach enabled resolution of action potentials as early as ∼8 ms post-stimulation without contamination by artifact.

Following spike sorting, we performed a per-trial approach implementing a multi-level artifact handling strategy to ensure balanced conditions across active and sham stimulation. Trials were initially flagged as outliers if their artifact end time exceeded the third quartile plus 1.5 times the interquartile range within their respective condition. To enforce strict trial matching, the index of each outlier was mapped to its relative position within a 50-trial block (e.g., trial 57 maps to position 7), and the union of these relative positions from both active and sham conditions created a definitive list of positions to exclude. All trials occupying these positions were systematically excluded from all blocks across both conditions, ensuring identical trial selection. For the remaining trials, spikes occurring within a per-channel artifact window (defined from -5 ms to 2 ms beyond the maximum artifact offset of the retained trials) were excluded from analysis. The same interval of exclusion was applied across conditions, with its offset determined by the latest artifact offset time across all retained trials from all conditions. Among 193 single units, 8 were excluded from analysis because their artifact offset exceeded 100 ms after stimulation, resulting in a total of 185 units included in the study. Among the 185 units included in the final analysis, a total of 29,550 active TMS trials were recorded. After excluding trials with artifacts, 25,771 trials (3779 excluded) were retained for analysis. For the sham TMS condition, a total of 18,456 trials were recorded across these same 185 units. After excluding trials with artifacts, 16,310 trials (2146 excluded) were retained for analysis.

### Single-unit spike detection, sorting, and quality assessment

Artifact-cleaned high-impedance microwire recordings were preprocessed in Matlab R2022a using demodulated band transform denoising methods. Prior to single-unit spike detection, the data were downsampled to 12 kHz and bandpass filtered between 200-4000 Hz. Spike detection and unit sorting employed a semi-automated workflow combining algorithmic classification with manual review. The computational approach utilized High Order Spectral Decomposition for feature extraction and waveform identification (Berger et al., 2026; Kovach et al., 2023; Kovach & Howard, 2019). This method estimated templates for putative action potentials on each channel through blind deconvolution techniques. Features derived from this process underwent clustering via Gaussian mixture modeling, and the resulting spike timestamps were used to generate candidate unit waveforms. Manual curation criteria included characteristic action potential morphology, waveform consistency across occurrence times within each cluster, and adherence to physiological absolute refractory period constraints (less than 1% of inter-spike intervals below 1 ms). The tendency for spikes to occur in brief high-frequency bursts (burst index) was quantified from the spike autocorrelogram using an established method (Royer et al., 2012). Autocorrelograms were computed with 1 ms bins and extended to 100 ms lag, then smoothed with a Gaussian kernel (σ=2 ms). The burst index was defined as the difference between the peak and baseline autocorrelogram values, normalized by the peak value. The peak was taken as the maximum value within the 0-10 ms lag window, and the baseline was defined as the mean value within the 40-50 ms lag window. Burst index values were constrained between -1 and 1, with positive values indicating burst-like firing and negative values indicating more regular firing patterns. The projection test was used to confirm distinct separation of each putative unit within a given bundle of 8 microwires. Temporal stability of units was ensured by visually confirming that each unit had spikes with consistent waveforms across the duration of the recording period. Single-unit quality metrics are provided in Supplementary Figure 2.

### Putative cell-type classification

Single units were classified into putative PYs and INs according to established methods in humans (Agopyan-Miu et al., 2023; Dickey et al., 2021; Kussovska et al., 2025; Peyrache et al., 2012). We performed unsupervised clustering using a Gaussian mixture model with *k*=2 clusters. The analysis was based on a feature matrix constructed from four electrophysiological parameters derived from each unit: trough-to-peak duration, full-width at half-maximum, burst index, and the log10-transformed firing rate. Prior to clustering, all features were *z*-scored to ensure equal weighting. The resulting two clusters were assigned PY or IN identities based on their mean trough-to-peak duration; the single units in the cluster exhibiting the shorter mean duration were classified as putative INs, consistent with their characteristic narrower spike waveforms, while the single units in the other cluster were classified as putative PYs.

### Intracranial TMS-evoked potentials

To compute iTEPs, intracranial macroelectrode recordings were epoched relative to the TMS pulse. To exclude channels contaminated by prolonged amplifier saturation, it was required that each channel’s average iTEP waveform have a zero-crossing within the 20-150 ms post-stimulus window for both active and sham conditions. For each trial, the iTEP waveform was baseline-corrected by subtracting the mean voltage from a pre-stimulus window (-550 to -50 ms) and then *z*-scored using the standard deviation of the same baseline period. The RMS of this processed signal was then calculated over the post-stimulus analysis window (50 to 500 ms) to yield a single iTEP magnitude value per trial for correlation with neuronal activity. High gamma amplitude was computed by bandpassing the broadband data from 70-190 Hz using a zero-phase shift 6th order Butterworth filter and then taking the amplitude of the Hilbert transform.

### Statistical methods

In order to identify significant stimulus-evoked changes in single-unit firing rates, we performed a trial-by-trial analysis for active and sham conditions. Single-unit spike times were binned into 1 ms intervals relative to the TMS pulse. For each trial, a baseline firing rate was calculated from the pre-stimulus period of -1010 ms to -10 ms. Post-stimulus firing rates were then calculated using a sliding window approach to detect significant changes from baseline. To detect rate increases, we used a 50 ms window advanced in 20 ms steps; for rate decreases, we used a 100 ms window advanced in 40 ms steps, with both analyses beginning after the artifact period and running up to 1000 ms post-stimulation. Within each window, the post-stimulation rate was compared to the trial-matched baseline rate using paired sign-flip one-sided permutation tests (1000 permutations), computed on the mean trial-wise rate difference and performed separately for increases and decreases. The resulting *p*-values were corrected for multiple comparisons using the False Discovery Rate (Benjamini & Hochberg, 1995) applied independently for each direction. A time point was considered to have a significant change if it belonged to a window that survived FDR correction at an alpha of 0.05 and was part of a sequence of at least two consecutive significant windows, with windows overlapping according to the step sizes. For visualization, peri-stimulus time histograms were generated using 20 ms bins and were also presented as z-scores, where the mean firing rate of each bin was normalized by subtracting the mean and dividing by the standard deviation of the collection of baseline firing rates from each individual trial.

To compare the proportion of units with significant firing rate modulation between striato-thalamic and limbic networks, we performed unit-level permutation tests treating each recording unit as an independent observation with a binary outcome (1=significant modulation, 0=no significant modulation). For units showing significant increases in firing rate, we tested the directional hypothesis that striato-thalamic neurons show a higher proportion of significant increases compared to limbic neurons using a one-sided permutation test. For units showing significant decreases in firing rate, we tested the directional hypothesis that limbic neurons show a higher proportion of significant decreases compared to striato-thalamic neurons using a one-sided permutation test. The permutation test procedure was as follows: (1) we calculated the observed difference in proportions between the two networks (striato-thalamic proportion - limbic proportion for increases, or limbic proportion - striato-thalamic proportion for decreases); (2) under the null hypothesis of no group difference, we randomly permuted group labels (striato-thalamic vs. limbic) 10,000 times while preserving the original sample sizes in each group; (3) for each permutation, we re-calculated the difference in proportions; (4) we computed the *p*-value by counting how many permuted differences were at least as large as the observed difference and dividing by the total number of permutations (10,000), adding one to both counts to prevent zero *p*-values. We performed the same such approach using a two-sided permutation test to assess for hemispheric differences of facilitation or suppression in striato-thalamic or limbic regions. This same permutation method was used to compare unit responses to active vs. sham stimulation.

To identify distinct response patterns, we performed unsupervised clustering on peri-stimulus time histograms from significantly modulated single units. Raw spike rate data were normalized using *z*-score transformation relative to a baseline period (-1010 to -10 ms), and a 50-600 ms post- stimulus window was extracted for clustering analysis after linearly interpolating across artifact periods and applying Gaussian smoothing (10-bin window with 20 ms per bin). We reduced data dimensionality using principal component analysis, selecting the minimum number of components required to capture at least 90% of variance (determined via scree analysis). Prior to clustering, outliers were identified and removed using an Isolation Forest algorithm with a contamination fraction of 0.05. We applied *k*-means clustering with 50 random initializations to ensure stable convergence, determining the optimal number of clusters (*k*) through a consensus approach integrating four metrics: Calinski-Harabasz index, Davies-Bouldin index, Gap statistic, and Silhouette coefficient, with the final *k* selected as the mode of the four optimal values (Huang et al., 2024). To assess whether network composition differed across identified clusters, we fit generalized linear mixed-effects models with binomial distribution for each network (striato-thalamic and limbic), using cluster assignment as a fixed effect and participant identity as a random effect to account for within-participant correlation of units, with statistical significance assessed via omnibus *F*-tests on fixed effects:

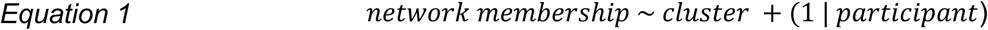

To compare the spike waveform and timing properties of PY and IN, we used linear mixed-effects (LME) models. For each unit characteristic (i.e., full-width at half-maximum, burst index, log-scale firing rate, and coefficient of variation 2 (CV2)), we modeled the metric as a function of the classified cell type as follows:

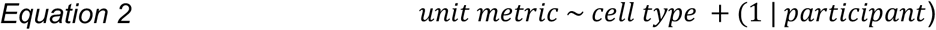

To control for the non-independence of neurons recorded from the same individual, the participant was included as a random intercept in the model. The statistical significance of cell type as a predictor for each metric was then determined from an ANOVA on the fitted LME model.

To compare the response characteristics of PY and IN, we performed statistical tests on both the proportion of responsive cells and the magnitude of their modulation. Differences in the proportion of PY vs. IN cells exhibiting specific response types (e.g., “increase only,” “decrease only”) were assessed using a one-sided permutation test (10,000 iterations) to evaluate directional hypotheses (e.g., that a greater proportion of INs increase their firing rate).

For the primary analyses of correlations between single-unit modulations and iTEPs (Figure 5), inclusion required each participant to have ≥5 macroelectrode bipolar re-referenced channels with iTEPs in each network (striato-thalamic and limbic) and ≥5 responsive and ≥5 non-responsive single units. Only participant 3 met all criteria (25 striato-thalamic macroelectrode channels, 6 limbic macroelectrode channels; 48 responsive units, 35 non-responsive units). Participants 1 and 4 were excluded due to insufficient macroelectrode coverage of one or both networks (participant 1: 0 striato-thalamic macroelectrode channels; participant 4: 4 limbic macroelectrode channels) and absent striato-thalamic responsive units (0 in each). For an exploratory supplementary analysis (Supplementary Figure 7B), we applied relaxed inclusion criteria requiring ≥5 macroelectrode channels in each network and ≥1 responsive single unit in at least one network. Under these criteria, participant 2 was additionally included (25 striato-thalamic macroelectrode channels, 8 limbic channels; 3 striato-thalamic responsive units, 12 limbic responsive units). Correlations were computed for each participant separately using the same statistical procedures described above.

To examine relationships between single-units modulations and iTEPs, we calculated Pearson correlations on a trial-by-trial basis between unit firing rates and iTEP root mean square (RMS) amplitudes during the analysis window (50-500 ms post-stimulus). Bipolar re-referenced macroelectrode channels were selected based on trans-gray matter placement with iTEP zero crossing within 20-150 ms after stimulation. For the primary analysis (participant 3: *n*=61 channels), correlations were computed separately for network-specific iTEPs (cortico-striato-thalamic and limbic networks) and separately for responsive (*n*=48) vs. non-responsive (*n*=35) single units during active and sham conditions. The same approach was applied to participant 2 (*n*=56 channels) in an exploratory supplementary analysis. Additionally, for network-specific analyses (Supplementary Figure 7), correlations were computed separately for single units classified as striato-thalamic or limbic, crossed with striato-thalamic and limbic network iTEPs. For each single unit, we assessed the correlation of the mean *z*-scored iTEP RMS (means computed separately for striato-thalamic and limbic networks) with the Poisson *z-*scored firing rate across trials (active TMS: *n*=150 trials per unit; sham TMS: *n*=100 trials per unit). Unit firing rates were Poisson *z*-scored relative to pre-stimulus baseline activity (-550 to -50 ms), while iTEP RMS amplitudes were *z*-scored relative to their pre-stimulus baseline (-550 to -50 ms). Poisson *z*-scoring yields a modulation score for each trial, where positive scores indicate activity exceeding the baseline expectation and negative scores indicate suppression. This approach adjusts for the relationship between the mean and variance inherent to Poisson-distributed count data, providing a more reliable measure of modulation than a standard Gaussian *z*-score when mean spontaneous spike counts on each trial are small. The baseline period *T*_base_, was used to estimate the mean spontaneous firing rate *λ*base, in units of spikes per second (Hz) across *N*_trials_:

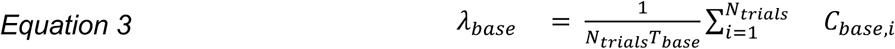

Where *C*_base,i_ is the number of spikes during the baseline window of trial *i*. Given the duration of the analysis window *T*_analysis_, the expected number of spikes (*µ*_expected_) and the expected standard deviation (*σ*_expected_) within that window are calculated assuming the Poisson distribution where the variance equals the mean:

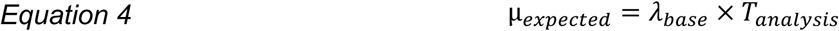

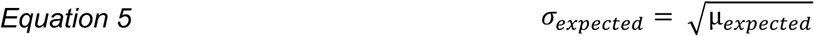

For each trial *i*, the Poisson *z*-score (*Z*_Poisson,i_) is then computed by comparing the observed spike count in the analysis window (*C*_analysis,i_) to the expected count, normalized by the expected standard deviation:

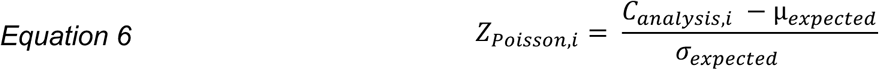

To control for multiple comparisons across single unit-iTEP correlation tests as well as across group contrasts, we applied FDR correction as described above. Correlation coefficients were Fisher *z*-transformed (*r_z_*) for normality prior to statistical evaluation of their distributions. Between-group comparisons of Fisher *z*-transformed correlation coefficients were performed using Wilcoxon rank-sum tests, while one-sample assessments of striato-thalamic and limbic network correlations utilized Wilcoxon signed-rank tests. For between-group comparisons of Fisher *z*-transformed correlation coefficients, one-sided Wilcoxon rank-sum tests were used to assess the directional hypothesis that correlations in the responsive-active group were significantly greater than those in the non-responsive and sham conditions.

## Acknowledgements

This work was supported by NIH R01DC004290 and NIH R01MH132074-01. The authors thank Benjamin Pace, Christopher Garcia, Danny Huang, Davide Momi, Ethan Solomon, Haiming Chen, Jade Truong, Kirill Nourski, Lily Forman, Nai-Feng Chen, Jessica Ross, Sabra Sisler, Samantha Gray, Sara Parmigiani, and Winn Hartford.

