## Supplementary Information for "Transcranial magnetic stimulation to the dorsolateral prefrontal cortex modulates single-neuron activity in humans"

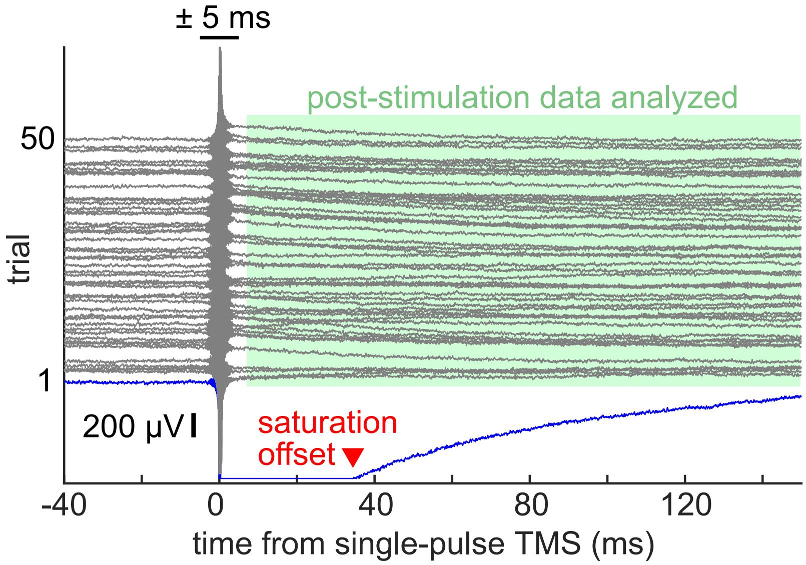


**Supplementary Figure 1: Artifact detection procedure.** Each trace shows an individual trial, with trials y-shifted for visualization purposes. The following steps were performed to address TMS artifacts in microwire recordings: (1) Each trial's artifact onset was set to -5 ms and offset was determined as the latest of +5 ms after single-pulse TMS or the amplifier saturation (example in red) or high-frequency offset time. (2) Trials with artifact offset times exceeding Q3 + 1.5 × IQR (e.g., the individual trial in blue) were identified as outliers and excluded across all conditions to ensure balanced trial selection. (3) Among the retained trials, artifacts were corrected from -5 ms to the artifact offset time (e.g., +5 ms in this example) using autoregressive interpolation (Cline et al., 2021). (4) Single-unit spikes were detected using the corrected trace. (5) Post-stimulation spiking was analyzed beginning at the maximum artifact offset across all retained trials plus 2 ms (e.g., +7 ms in this example, indicated in green), with this uniform exclusion window applied across all conditions. IQR, inter-quartile range; Q3, third quartile; TMS, transcranial magnetic stimulation.

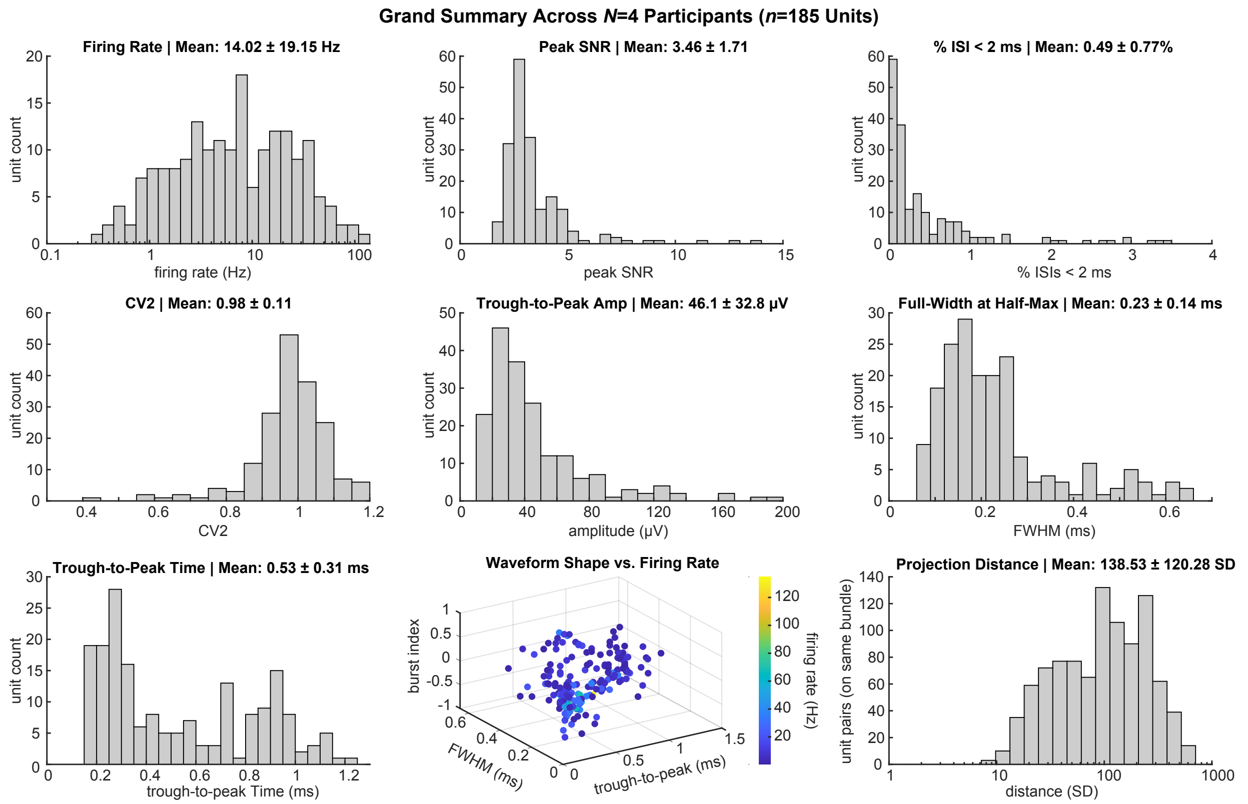


**Supplementary Figure 2: Single-unit quality metrics.** Histograms of single-unit quality metrics indicate large SNR, minimal contamination of spikes within 2 ms, and clear separation (isolation distance). Mean and standard deviation displayed in each plot title. CV, coefficient of variation 2; FWHM, full-width at half-maximum; ISI, inter-spike interval; SD, standard deviation; SNR, signal-to-noise ratio.

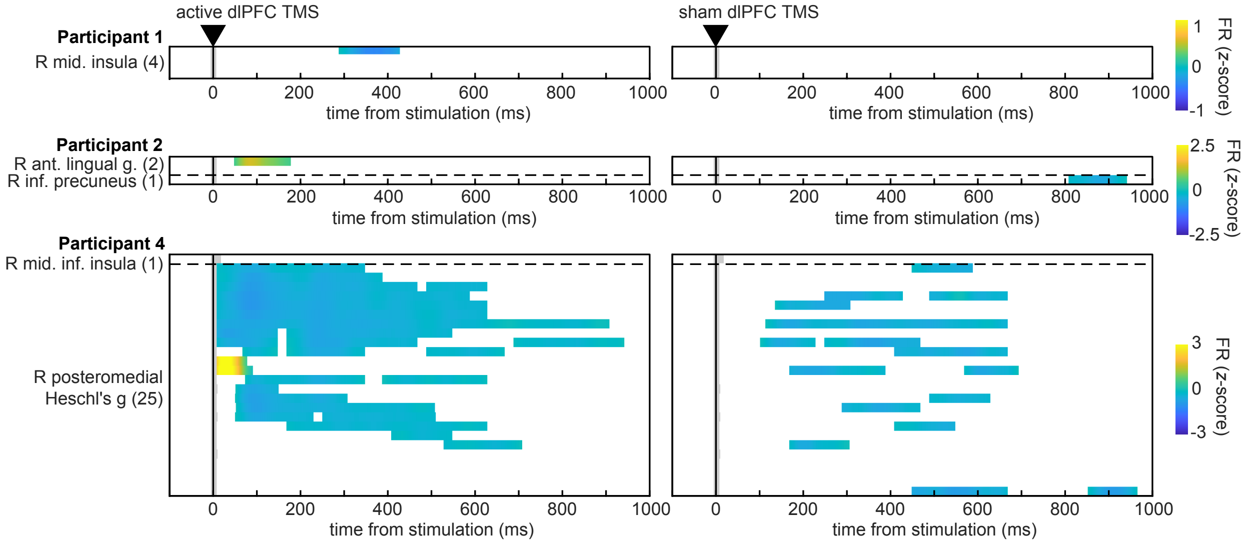


**Supplementary Figure 3: Single-unit responses in non-striato-thalamic and non-limbic areas.** Heatmaps show *z*-scored firing rates across single units from non-striato-thalamic and non-limbic regions (*n*=33 units, *N*=3 participants; participant 3 had no single units outside striato-thalamic and limbic regions) during active dlPFC TMS (left) and sham dlPFC TMS (right). Each row represents one unit, with only statistically significant time bins displayed in color. Regions are separated by dashed lines. Within each region, units are sorted by the latency of their first significant firing rate change. Numbers in parentheses indicate the total units recorded in each region from each participant. See Figure 2B for single-unit responses from striato-thalamic and limbic single units. FR, firing rate.

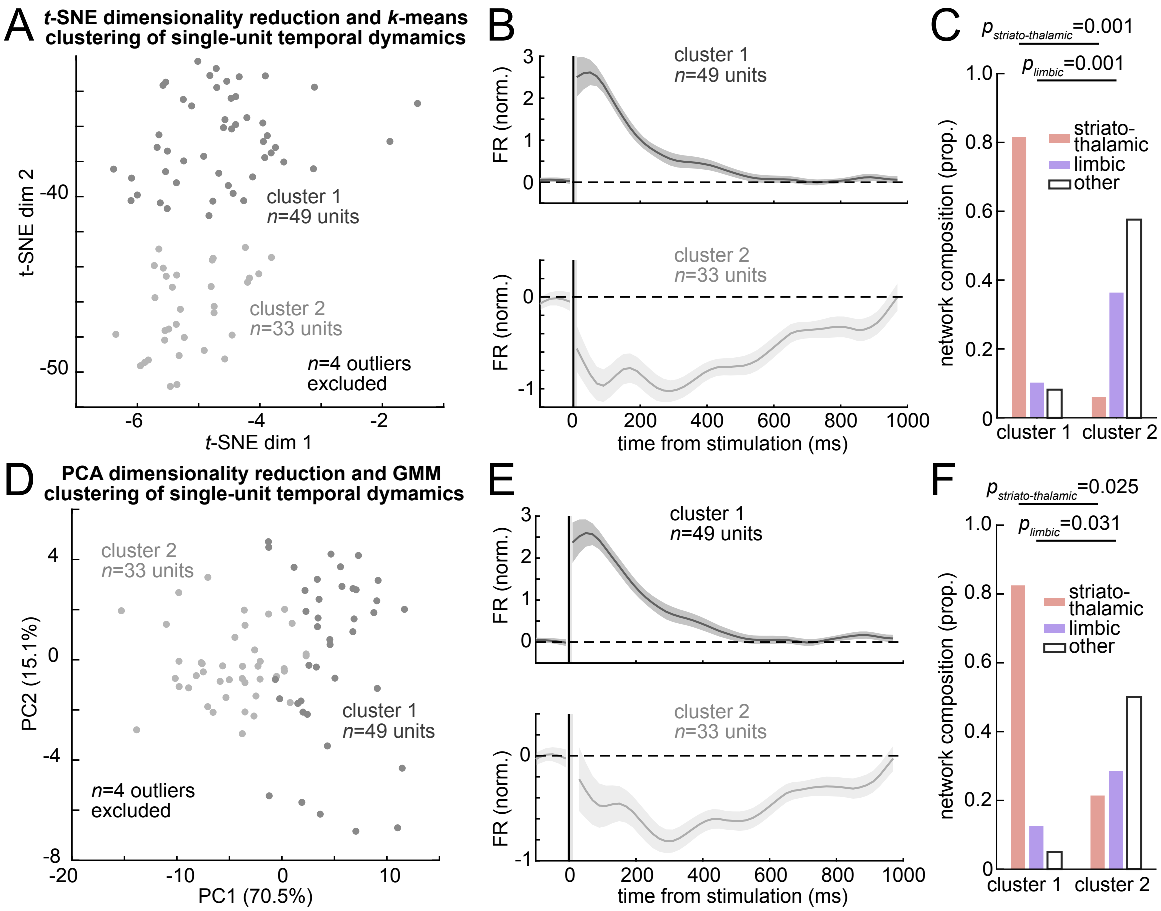


**Supplementary Figure 4: Alternative dimensionality reduction and clustering methods confirm two distinct single-unit response phenotypes.** While Figure 3D-E utilized PCA for dimensionality reduction and *k*-means for clustering single-unit response dynamics, the following alternative methods reveal the same two distinct response phenotypes: one with rapid facilitation localizing to striato-thalamic areas and another with slower suppression localizing to limbic areas. (A) *t*-SNE embedding of normalized firing rate responses across all significantly modulated single units (*n*=82 units; *n*=4 outliers excluded). *k*-means clustering identified two distinct temporal response patterns: cluster 1 (*n*=49 units) and cluster 2 (*n*=33 units). (B) Mean and SEM normalized firing rates for each cluster in A. Top shows cluster 1, which exhibits facilitation peaking at ~80 ms and decaying until ~600 ms. Bottom shows cluster 2, which exhibits suppression peaking at ~300 ms and sustained until ~1000 ms. (C) Quantification of network composition (striato-thalamic, limbic, or other) by cluster reveals that cluster 1 with early peaking facilitation contains significantly more striato-thalamic single units ($p$=0.001) whereas cluster 2 with sustained suppression contains significantly more limbic single units ($p$=0.001). (D-F) Same as A-C except using PCA for dimensionality reduction and Gaussian mixture model (GMM) clustering. GMM identified cluster 1 (*n*=49 units) and cluster 2 (*n*=33 units) with similar network composition differences ($p_{striato\text{-}thalamic}$=0.025, $p_{limbic}$=0.031). PCA, principal component analysis; *t*-distributed Stochastic Neighbor Embedding.

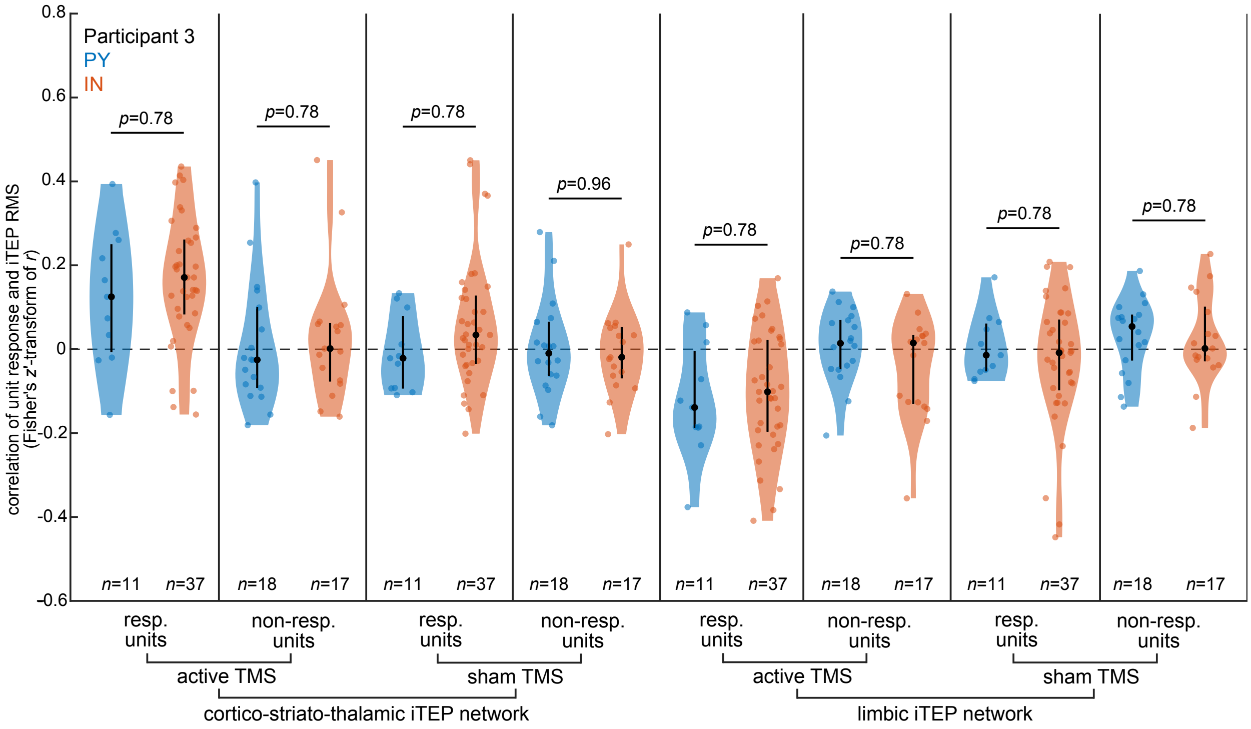


**Supplementary Figure 5: Single-unit correlations with network iTEPs do not differ between putative pyramidal cells and interneurons.** Same format as Figure 5G, with each distribution split by putative cell type: pyramidal cells (PY, blue) and interneurons (IN, orange). Correlations between single-unit response and cortico-striato-thalamic iTEP RMS (left four violin pairs) and limbic network iTEP RMS (right four violin pairs) are shown for responsive and non-responsive units during active and sham TMS in participant 3. No significant differences between PY and IN were observed in any condition (two-sided Wilcoxon rank-sum tests; all $p_{FDR} \geq 0.78$). Black circles show median and vertical bars show interquartile range. RMS, root mean square.

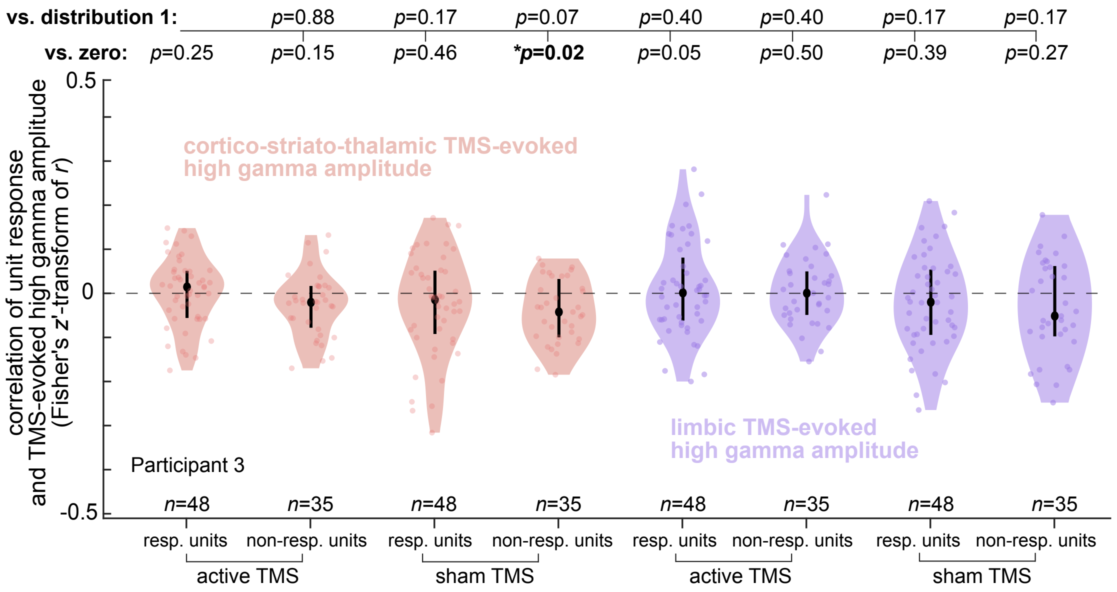


**Supplementary Figure 6: Single-unit responses are not robustly correlated with TMS-evoked high gamma amplitude.** Same format as Figure 5G, with trial-by-trial correlations (Fisher's *z*-transformed) computed between single-unit firing rate modulation and TMS-evoked high gamma (70-190 Hz) amplitude rather than iTEP RMS. Data are shown for responsive and non-responsive units during active and sham TMS in participant 3, with cortico-striato-thalamic high gamma amplitude (red, left four violins) and limbic high gamma amplitude (purple, right four violins). FDR-corrected *p*-values from two-sided Wilcoxon signed-rank tests against zero are shown above each violin (vs. zero), and FDR-corrected *p*-values from two-sided Wilcoxon rank-sum tests comparing each distribution to responsive units during active TMS with cortico-striato-thalamic high gamma (distribution 1) are shown at top (vs. distribution 1). While non-responsive units during sham TMS with cortico-striato-thalamic high gamma reached significance against zero ($p_{FDR}$=0.02) this was not significantly different from distribution 1 ($p_{FDR}$=0.07), and no other conditions were significant (all $p_{FDR} \geq 0.05$ vs. zero; all $p_{FDR} \geq 0.17$ vs. distribution 1). Black circles show median and vertical bars show interquartile range. *p<0.05.

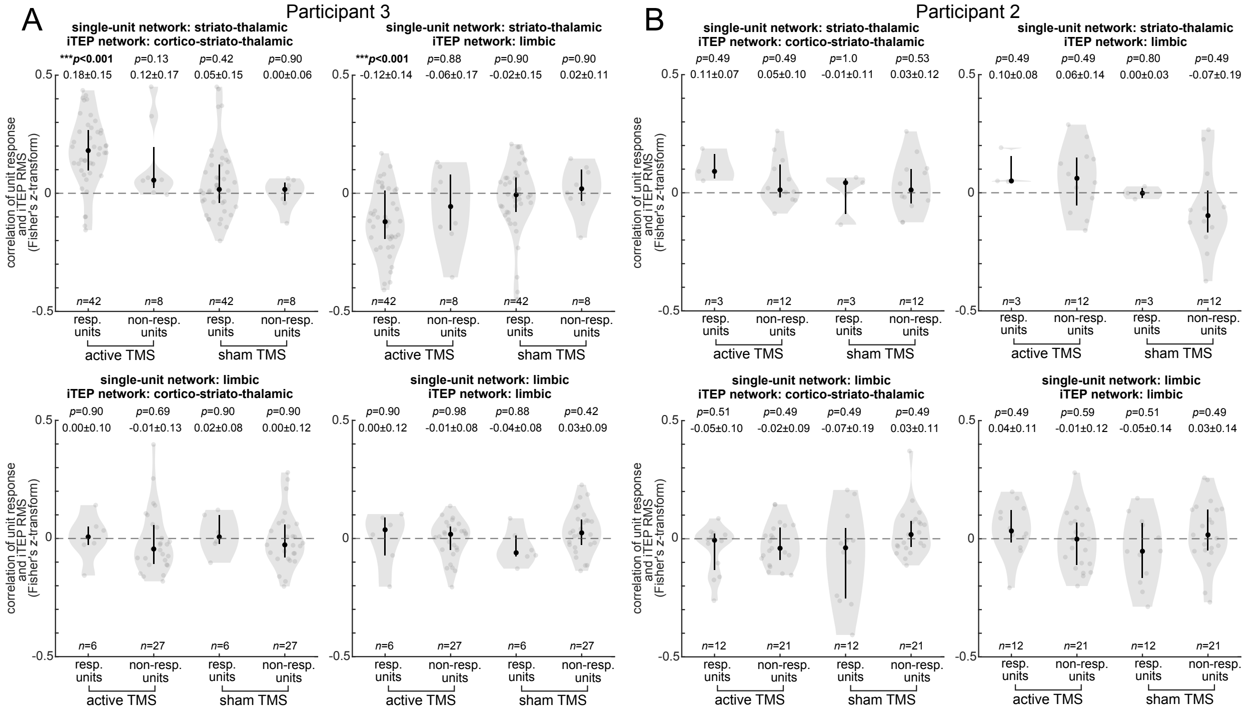


**Supplementary Figure 7: Striato-thalamic and limbic single-unit relationships with cortico-striato-thalamic and limbic iTEPs across two participants.** Trial-by-trial correlations (Fisher's *z*-transformed) between single-unit firing rate modulation and iTEP RMS amplitude, organized by single-unit region (rows: striato-thalamic, limbic) and macroelectrode network iTEP (columns: cortico-striato-thalamic, limbic). Violin plots show correlations for responsive units during active TMS, non-responsive units during active TMS, responsive units during sham TMS, and non-responsive units during sham TMS. Black circles show median and vertical bars show interquartile range. Mean and standard deviation as well as within-participant FDR-corrected *p*-values from two-sided Wilcoxon signed-rank tests against zero are shown above each distribution. (A) Participant 3 (primary analysis in Figure 5). Responsive striato-thalamic single units to active TMS were positively correlated with cortico-striato-thalamic iTEPs (mean $r_z$=0.18±0.15, $p_{FDR}$<0.001; top left) and anti-correlated with limbic network iTEPs (mean $r_z$=-0.12±0.14, $p_{FDR}$<0.001; top right). No significant correlations were observed for limbic single units with either network (bottom row; all $p_{FDR}$≥0.42), for non-responsive units, or during sham TMS (all $p_{FDR}$≥0.13). (B) Participant 2 (exploratory analysis) had adequate macroelectrode coverage of both networks (*n*=25 striato-thalamic and *n*=8 limbic channels) but a limited number of responsive striato-thalamic single units (*n*=3). Responsive striato-thalamic single units to active TMS showed a positive but non-significant mean correlation with cortico-striato-thalamic iTEPs (mean $r_z$=0.11±0.07, $p_{\mathrm{FDR}}$=0.49; top left). No significant anti-correlation with limbic network iTEPs was observed (mean $r_z$=0.10±0.08, $\mathrm{p_{FDR}}$=0.49; top right). Responsive limbic single units (*n*=12) to active TMS showed no significant correlation with either cortico-striato-thalamic (mean $r_z$=-0.05±0.10, $p_{FDR}$=0.51) or limbic network iTEPs (mean $r_z$=0.04±0.11, $p_{\mathrm{FDR}}$=0.49). No other correlations reached significance (all $p_{\mathrm{FDR}}$=0.49-1.0). The limited number of responsive striato-thalamic units in this participant substantially constrained statistical power, precluding definitive conclusions regarding the generalizability of the correlations observed in participant 3.

| Partic-ipant | Age | Sex | Handed-ness | Seizure onset zone | ASM | Mood status and psychiatric history | Microwire regions implanted | Macro-electrode regions | Days post-implant | TMS target | No. TMS pulses | ISI | TMS intensity |
|---|---|---|---|---|---|---|---|---|---|---|---|---|---|
| 1 | 35 | F | R | R amygdala/ hippocampus; R lingual gyrus; R posterior cingulate; L hippocampus | cenobamate, topiramate, levetiracetam, lamotrigine, gabapentin | no mood episode during study; history of unspecified psychosis (none during study), tobacco use disorder, and claustrophobia | R anterior insula, R middle insula, R post hippocampus, R medial occipital, R parahippocampal gyrus, L amygdala, L middle hippocampus, L posterior hippocampus | N/A | 13 | L dlPFC | 50 sham, 50 sham TAAC, 50 active, 50 active TAAC, 50 sham, 50 sham TAAC, 50 active, 50 active TAAC | 3 s | 100% rMT |
| 2 | 42 | F | R | R amygdala, R middle hippocampus, R posterior hippocampus | clonazepam, divalproex, midazolam (rescue) | no mood episode during study; history of recurrent MDD and GAD | R subgenual cingulate, R anterior cingulate, R middle cingulate, R posterior cingulate, R putamen, R amygdala, R middle hippocampus, R posterior hippocampus, R thalamus, R inferior precuneus, R anterior lingual gyrus | angular gyrus, Heschl's gyrus, posterior insula, inferior frontal gyrus, middle frontal gyrus, supramarginal gyrus, anterior insula, anterior cingulate, middle anterior cingulate, orbital gyrus, gyrus rectus, planum polare, middle temporal gyrus, superior temporal sulcus, planum temporale, precentral gyrus, precuneus, superior temporal gyrus, lingual gyrus | 19 | L dlPFC | 50 sham TAAC, 50 sham, 50 active TAAC, 50 active, 50 active | 2.5 s | 120% rMT |
| 3 | 47 | M | R | L insula | zonisamide, lamotrigine | major depressive episode during study; history of recurrent MDD | R amygdala, R thalamus x 2, L amygdala, L middle hippocampus, L posterior hippocampus, L putamen x 4, L pulvinar | inferior frontal g, parietal operculum, posterior insula, postcentral gyrus, putamen, pulvinar, supramarginal gyrus, putamen, amygdala, anterior insula, planum polare, Heschl's gyrus, middle temporal gyrus, superior temporal gyrus, planum temporale | 15 | L dlPFC | 50 sham, 50 sham, 50 active TAAC, 50 active, 50 active TAAC | 2.5 s | 120% rMT |
| 4 | 56 | F | R | R superior/ middle frontal | clobazam, eslicarbazepine, clonazepam, gabapentin, levetiracetam, midazolam (rescue only) | no mood episode during study; no psychiatric history | R subgenual cingulate, R post superior insula, R anterior inferior insula, R middle inferior insula, R posteromedial Heschl's g, L subgenual cingulate, L middle cingulate | N/A | 12 | L dlPFC | 50 sham TAAC e-stim, 50 sham, 50 active TAAC, 50 active TAAC, 50 active TAAC, 50 active TAAC, 50 active TAAC | 2.5 s | 120% rMT |

**Supplementary Table 1: Participant demographics, recording characteristics, and stimulation paradigms.** ASM=anti-seizure medication used during inpatient stay, dlPFC=dorsolateral prefrontal cortex, E-stim=scalp electrical stimulation, GAD=generalized anxiety disorder, ISI=inter-stimulus interval, MDD=major depressive disorder, rMT=resting motor threshold, TAAC=TMS Adaptable Auditory Control, TMS=transcranial magnetic stimulation.

| Network | Parcellations |
|---|---|
| cortico-striato-thalamic | putamen, thalamus, posterior insula, parietal operculum, supramarginal gyrus, angular gyrus, middle frontal gyrus, inferior frontal gyrus |
| limbic | amygdala, hippocampus, parahippocampal gyrus, cingulate cortex, anterior insula, orbital gyrus |

**Supplementary Table 2: Cortico-striato-thalamic and limbic anatomical parcellations.** Only regions sampled by microwire or macroelectrode recordings which were included in the study are listed. Among cortico-striato-thalamic areas, microelectrodes were only implanted in striatal and thalamic regions. See Supplementary Table 1 for participant-specific data.
